# Enzymado: Are Protein Language Models Getting Better at Enzyme Tasks?

**DOI:** 10.64898/2026.11.26.769339

**Authors:** Jose Manuel Barraza-Chavez, Ricardo Almada-Monter, William Chau, Rana A. Barghout, Aziz Abdur Rakib, Maxim Kirby, Radhakrishnan Mahadevan, Adrian Jinich, Benjamin Sanchez-Lengeling

**Affiliations:** ChemE, University of Toronto, UC San Diego; Vector Institute, UC San Diego; Chem. & Biochem., UC San Diego, UC San Diego; Biochem. & Mol. Biophys., UC San Diego

## Abstract

Protein language model (PLM) embeddings are a common representation for enzyme engineering tasks. PLMs are pretrained using vast protein data, and while enzymes are proteins, they are distinguished by their catalytic nature, which is dependent on substrates and reaction mechanisms. Current PLM evaluations test isolated properties with incompatible data, splits, and metrics, leaving open how PLMs can better serve enzymes and what drives their performance. We present Enzymado, a framework for evaluating enzyme representations, with property datasets curated from public databases and experimental protein engineering studies, spanning kinetics, mutation effects, function, binding, solubility, and residue annotations. Enzymado standardizes data cleaning, evaluation splits, prediction heads, and metrics throughout the comparison of 14 PLMs. Representations compete through paired-bootstrap matches with ties set by measured training-seed noise, and a weighted Bradley–Terry model aggregates these matches into ratings with confidence intervals. Datasets, models, and properties are presented as executable cards, so that rating differences trace to modeling attributes and become testable hypotheses. We find that newer PLMs do not rank higher: the six released after December 2024 have the six lowest ratings. Neither parameter count nor embedding size predicts the best rating, while structure-aware pretraining leads, most clearly on catalytic residue tasks. The predictor head has a comparable effect to PLM choice on ratings, and rankings transfer only partly to distant homologs and engineering campaigns. New datasets enter through a documented ingestion workflow, which an LLM coding agent followed to add 28 of the 36 case studies; code, data, and cards are publicly available on GitHub with a live leaderboard offering practical modeling recommendations.

## 1 Introduction

Enzymes catalyze most chemical reactions in biology, making them both common targets for pharmaceuticals and tools for replacing industrial chemical processes [9, 19, 33, 62]. Current industrial applications of enzymes range from anti-diabetic drug synthesis to stain removers in detergents and PET plastics degradation [22,41,60]. Enzyme properties, such as substrate specificity, kinetics, and stability, play an important role in these discovery and engineering campaigns. However, large-scale measurement of different properties across different conditions remains a bottleneck. Therefore, computational methods that predict enzyme properties from enzyme sequences are increasingly used to narrow the experimental space of enzyme exploration [51, 81, 84]. Most of these methods use protein language models (PLMs) to represent enzyme sequences as numerical vectors that are then used for downstream prediction.

PLMs such as ESM3 [30], Ankh [20], and ProstT5 [32] vary in training data, training objectives, and architecture, leading to diverse protein representations. These PLMs are trained on millions of protein sequences, and their representations have been shown to encode biologically relevant information such as evolutionary relationships [23], contacts [57, 71], secondary structure and subcellular localization [31], and function [49, 80]. ESM3 uses a bidirectional transformer that was trained to predict masked information across sequence, structure, and functional annotations. Ankh uses an encoder–decoder transformer trained on UniRef50 sequences to reconstruct masked sequence content [20]. ProstT5 builds on the encoder–decoder architecture and pretrained weights of ProtT5, with additional training on amino acid sequences paired with structural representations derived from AlphaFold predictions. Its training includes bidirectional translation between amino acid sequences and 3D tokens, which encode three-dimensional interactions between residues.

Despite the progression and variety of PLMs, recent studies have discovered limitations in the performance of the representations across different enzyme property tasks. Li *et al.* found that improved pretraining performance does not consistently translate into better downstream predictions across protein tasks such as thermostability and mutational effects [46]. For enzyme function annotation, de Crécy-Lagard *et al.* documented incorrect EC assignments and limited success in making novel functional predictions for uncharacterized proteins [18]. For enzyme kinetics, Kroll *et al.* showed that DLKcat predictions deteriorate for enzymes different from those in the training set, whereas their PLM-based TurNuP model improves generalization in this setting [43]. Wang *et al.* further found that ProtT5 representations outperform ESM2 representations for turnover-number prediction in their evaluation, and that random data splits can overestimate kinetic-prediction performance compared with sequence-clustered evaluation [72]. Selecting a PLM sets the foundation for downstream modeling tasks and strongly determines predictive performance, yet holistic evaluations of what PLMs do best across different tasks are rare.

This motivates the evaluation of protein representations across enzyme properties under consistent conditions to explicitly assess a representation’s generalization and understanding of enzymes. From this, a practical question arises: *how do we know which PLM representation to choose?* A good representation can be defined as one that encodes the necessary information such that even a simple downstream predictive model can use it and perform well [10]. Since these representations come from the latent space of PLMs, determining the information encoded by the representations is not a trivial task [7]. More questions arise as new PLMs are developed: *How do we build better representations for enzymes? How much better are the new representations? And which protein representations are better for enzyme tasks? (Fig. 1)*

**Figure 1:**
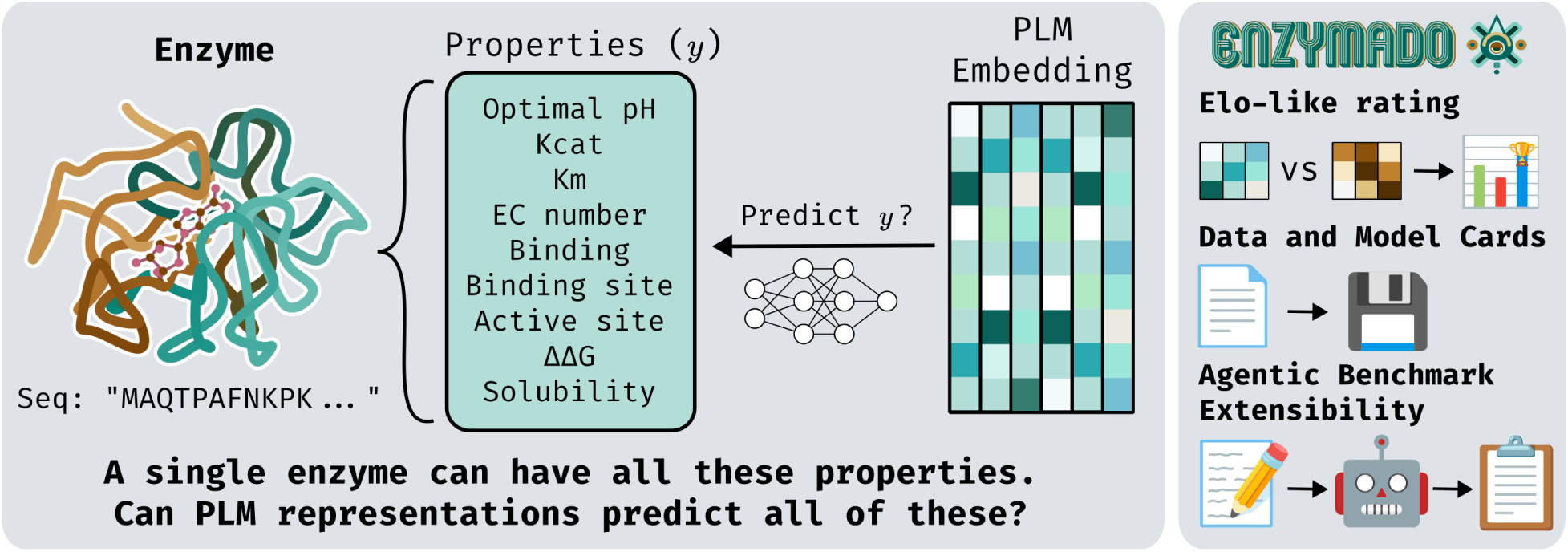
Enzymado probes whether PLM representations encode information for multi-property enzyme tasks. An enzyme can be defined by many properties, and Enzymado tests whether PLMs provide the necessary information for multi-property prediction.

A common way to approach this last question is to evaluate the predictive performance of different PLMs and models under the same tasks using benchmarks. These benchmarks often evaluate a specific and narrow set of properties. In addition, the benchmarks are fragmented across tasks, data cleaning processes, evaluation methods, and metrics. For example, ProteinGym [55], PEER [76], and CASP [82] cover mutation effects, general protein tasks, and structure, while CatPred-DB [11] and FireProtDB [64] cover kinetic parameters and stability. Most datasets and benchmarks cover different types of proteins indiscriminately, and do not evaluate the performance of PLMs across enzymes specifically. Enzymes have qualities that separate them from other proteins, such as their ability to catalyze reactions, which is a property defined not only by the enzyme but also by the substrate and the conditions. Enzyme-specific benchmarks such as CARE [77] focus on EC-number prediction and reaction-based retrieval. These tasks evaluate enzyme function, but do not show how the same representations perform on other enzyme properties, such as kinetics and stability.

To address this gap, we introduce **Enzymado**, a framework for evaluating protein representations across enzyme tasks (Fig. 2). We evaluate 14 PLM representations paired with five predictor architectures across 46 datasets (4,200 trained models, plus 3,844 seed runs) to test what biological and mechanistic information is linearly or nonlinearly recoverable from frozen sequence embeddings. The main contributions of this work are:

- **A noise-calibrated statistical rating methodology.** To aggregate incomparable metrics across classification and regression tasks without conflating optimization noise with model superiority, we formulate a weighted Bradley–Terry model with tie thresholds calibrated to empirical training-seed variance over 1,000 paired-bootstrap matches (Section 2.5, Appendix E).
- **Representation learning findings at scale.** Across 4,200 main training runs, we demonstrate that newer and larger PLMs do not yield superior enzyme representations (*ρ* = 0.58 with release date), that structure-aware pretraining drives performance on catalytic residue tasks, and that catalytic site information is encoded nonlinearly, causing linear probes to misrank structure-trained PLMs (Section 3).
- **Biologically faithful task formulations and multi-regime splits.** We standardize 10 database-derived property tasks and 36 experimental protein engineering campaigns across random, sequence-similarity, and stratified splits. Tasks are modeled to reflect their underlying assay structure, including substrate- and condition-aware encodings, mutant-to-wild-type difference representations, and hierarchical multi-label EC prediction with partial-level loss masking (Section 2.1, Section 2.2).
- **Data-centric cards and agentic ingestion.** We introduce model and dataset cards to structure prior knowledge (with 1,895 quotes verified against source pages), enabling us to test hypotheses across models (e.g., why model A beats B) and recommend models for specific enzyme systems. For continuous updates, our ingestion workflow adds datasets without invalidating stored embeddings; this agentic workflow was used to add 28 of the 36 case studies and surface source data errors (Section 2.6, Appendix M).

**Figure 2:**
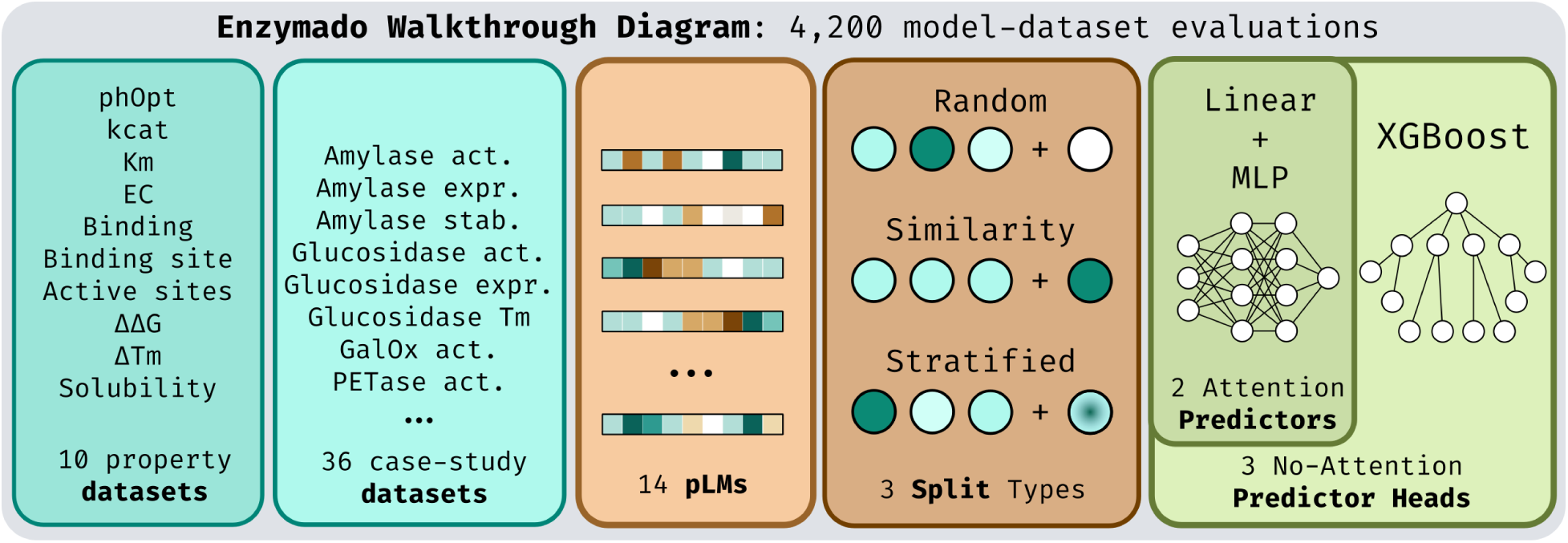
Enzymado walkthrough diagram and design choices. 10 property datasets plus 36 case-study datasets, across 14 PLMs, and 3 different split strategies, random and similarity for the property suite, and stratified for the case-study suite. 5 different predictor heads, linear and MLP heads with attention plus their no-attention variants and XGBoost.

## 2 Enzymado Design and Methods

### 2.1 Tasks and Data Sources

Enzymado contains 10 database-derived property datasets, comprising 5 regression and 5 classification tasks (Table 1), and 36 case-study datasets (Appendix Table 2). The property suite covers enzyme kinetics, optimal pH, mutation-induced stability changes, protein–ligand binding, EC annotation, residue annotation, and solubility. Each dataset is restricted to enzymes using the source’s enzyme annotations or a match to an EC-annotated Swiss-Prot sequence. Solubility uses the 3,664 enzyme sequences retained from the 71,421-sequence DeepSol dataset as distributed with Ankh [20, 40]. The sources include CatPred-DB [11], BRENDA [61], FireProtDB [64], BioLiP [78, 83], and UniProt/Swiss-Prot [6, 66, 67]. These tasks differ in their observational unit and in what a label means; Table 1 summarizes their sources and sizes, while Appendix I defines the labels and metrics.

**Table 1:** The 10 property datasets after their cleaning: rows, distinct sequences, and minimum, median, and maximum length over distinct sequences. Counts reflect the filters used in the reported runs, including fragment filtering where applicable and shared embedding coverage. *Scope*: global for the whole protein, local for residues.

| **Property** | **Type** | **Scope** | **Source** | **Rows** | **Proteins** | **Lengths (aa)** |  |  |
|---|---|---|---|---|---|---|---|---|
|  |  |  |  |  |  | **Min.** | **Median** | **Max.** |
| *k*<sub>cat</sub> | reg. | global | CatPred-DB | 22,693 | 7,012 | 10 | 385 | 2,515 |
| *K*<sub>M</sub> | reg. | global | CatPred-DB | 40,088 | 12,004 | 10 | 399 | 2,590 |
| pH<sub>opt</sub> | reg. | global | BRENDA | 11,113 | 10,770 | 39 | 411 | 2,836 |
| Δ*T*<sub>m</sub> | reg. | global | FireProtDB | 3,338 | 2,312 | 96 | 231 | 2,527 |
| ΔΔ*G* | reg. | global | FireProtDB | 3,760 | 2,518 | 96 | 164 | 1,863 |
| binding | cls. | global | BioLiP | 28,316 | 11,233 | 31 | 337 | 2,652 |
| EC number | cls. | global | Swiss-Prot | 233,586 | 233,534 | 30 | 360 | 2,860 |
| active sites | cls. | local | Swiss-Prot | 89,329 | 89,329 | 8 | 351 | 2,799 |
| binding sites | cls. | local | Swiss-Prot | 157,124 | 157,124 | 8 | 369 | 2,860 |
| solubility | cls. | global | DeepSol | 3,664 | 3,664 | 61 | 330 | 1,645 |

**Table 2:** The 36 case studies. *Layout*: A for protein only, B for protein and substrate (Table 15). *Rows* are the published rows after cleaning, *Seq.* the distinct sequences, and *Sub.* the distinct substrates, or condition combinations for the PET hydrolases, with n/a where the dataset has neither. *^†^* marks the 14 datasets whose sequences repeat across rows. Observables are abbreviated; each dataset card gives the exact definition and units.

| Dataset | Layout | Observable | Rows | Seq. | Sub. | Ref. |
|---|---|---|---|---|---|---|
| *Variant libraries of one parent enzyme* |  |  |  |  |  |  |
| align_amylase_activity | A | specific activity | 9,031 | 9,031 | n/a | [5] |
| align_amylase_expression | A | secreted expression | 9,421 | 9,421 | n/a | [5] |
| align_amylase_stability | A | thermostability | 8,736 | 8,736 | n/a | [5] |
| align_glucosidase_activity | A | activity | 354 | 354 | n/a | [5] |
| align_glucosidase_expression | A | expression | 324 | 324 | n/a | [5] |
| align_glucosidase_melting | A | melting point | 106 | 106 | n/a | [5] |
| amie_acetamide_dms | A | growth fitness | 6,290 | 6,290 | n/a | [73] |
| amie_isobutyramide_dms | A | growth fitness | 6,229 | 6,229 | n/a | [73] |
| amie_propionamide_dms | A | growth fitness | 6,292 | 6,292 | n/a | [73] |
| cyp2c9_abundance | A | abundance score | 6,371 | 6,371 | n/a | [3] |
| cyp2c9_activity | A | activity score | 6,143 | 6,143 | n/a | [3] |
| daox_epseq_activity | A | activity fitness | 6,404 | 6,404 | n/a | [70] |
| daox_epseq_stability | A | display expression | 6,434 | 6,434 | n/a | [70] |
| goase_variant_substrate<sup>†</sup> | B | oxidation activity | 1,183 | 34 | 54 | [65] |
| mcba_reaction_activity<sup>†</sup> | B | amidation activity | 10,918 | 1,217 | 9 | [45] |
| mpro_fret_cleavage | A | cleavage score | 5,745 | 5,745 | n/a | [25] |
| pafa_htmek_kcat | A | *k*<sub>cat</sub> | 1,018 | 1,018 | n/a | [52] |
| pafa_htmek_kcat_km | A | *k*<sub>cat</sub>/*K*<sub>M</sub> | 1,042 | 1,042 | n/a | [52] |
| pafa_htmek_km | A | *K*<sub>M</sub> | 1,018 | 1,018 | n/a | [52] |
| plpro_abundance | A | abundance fitness | 5,335 | 5,335 | n/a | [75] |
| plpro_activity | A | activity fitness | 5,335 | 5,335 | n/a | [75] |
| pyks_solubility | A | solubility score | 6,486 | 6,486 | n/a | [74] |
| trpb3_epistasis | A | growth fitness | 7,970 | 7,970 | n/a | [37] |
| welo5_soraphen_activity | A | chlorination conversion | 504 | 504 | n/a | [13] |
| *Panels of different enzymes* |  |  |  |  |  |  |
| ao_3fcr_transaminase<sup>†</sup> | B | specific activity | 2,034 | 79 | 30 | [4] |
| cleveland_galox<sup>†</sup> | B | specific activity | 385 | 12 | 40 | [16] |
| erickson_petase<sup>†</sup> | A | product release | 570 | 19 | 30 | [22] |
| fisher_fdh_nabr<sup>†</sup> | B | bromination conversion | 2,604 | 42 | 62 | [24,29] |
| fisher_fdh_nacl<sup>†</sup> | B | chlorination conversion | 2,604 | 42 | 62 | [24,29] |
| goldman_aminotransferase<sup>†</sup> | B | specific activity | 432 | 24 | 18 | [29,47] |
| goldman_olea_thiolase<sup>†</sup> | B | hydrolysis activity | 550 | 73 | 14 | [29,58] |
| huang_had_phosphatase_c0<sup>†</sup> | B | phosphate release | 5,610 | 34 | 165 | [29,35] |
| huang_had_phosphatase_c1<sup>†</sup> | B | phosphate release | 15,840 | 96 | 165 | [29,35] |
| huang_had_phosphatase_c2<sup>†</sup> | B | phosphate release | 7,920 | 48 | 165 | [29,35] |
| koudelakova_hld_panel<sup>†</sup> | B | specific activity | 390 | 13 | 30 | [42] |
| martinez_esterase_panel<sup>†</sup> | B | specific activity | 14,016 | 146 | 96 | [29,53] |

**Table 3:** The 24 variant-library datasets. All sequences within a dataset have the same length. Substitution counts are Hamming distances to a reference formed by the consensus residue at each position; this reference is not necessarily the experimental parent or wild type. A minimum of 0 means that the consensus reference occurs among the rows. *Pos.* counts the positions that vary anywhere in the dataset.

| **Dataset** | **Length (aa)** | **Substitutions** |  |  | **Pos.** |
|---|---|---|---|---|---|
|  |  | **Min.** | **Median** | **Max.** |  |
| align_amylase_activity | 425 | 0 | 1 | 11 | 425 |
| align_amylase_expression | 425 | 0 | 1 | 11 | 425 |
| align_amylase_stability | 425 | 0 | 1 | 11 | 425 |
| align_glucosidase_activity | 445 | 0 | 1 | 1 | 150 |
| align_glucosidase_expression | 445 | 0 | 1 | 1 | 136 |
| align_glucosidase_melting | 445 | 0 | 1 | 1 | 68 |
| amie_acetamide_dms | 346 | 0 | 1 | 1 | 341 |
| amie_isobutyramide_dms | 346 | 0 | 1 | 1 | 341 |
| amie_propionamide_dms | 346 | 0 | 1 | 1 | 341 |
| cyp2c9_abundance | 490 | 0 | 1 | 1 | 486 |
| cyp2c9_activity | 490 | 0 | 1 | 1 | 477 |
| daox_epseq_activity | 365 | 1 | 1 | 1 | 364 |
| daox_epseq_stability | 365 | 1 | 1 | 1 | 364 |
| goase_variant_substrate | 639 | 0 | 2 | 3 | 11 |
| mcba_reaction_activity | 518 | 0 | 1 | 1 | 64 |
| mpro_fret_cleavage | 306 | 0 | 1 | 1 | 303 |
| pafa_htmek_kcat | 529 | 0 | 1 | 2 | 526 |
| pafa_htmek_kcat_km | 529 | 0 | 1 | 2 | 526 |
| pafa_htmek_km | 529 | 0 | 1 | 2 | 526 |
| plpro_abundance | 315 | 1 | 1 | 1 | 315 |
| plpro_activity | 315 | 1 | 1 | 1 | 315 |
| pyks_solubility | 439 | 0 | 1 | 1 | 439 |
| trpb3_epistasis | 397 | 0 | 3 | 3 | 3 |
| welo5_soraphen_activity | 290 | 1 | 3 | 3 | 3 |

For BioLiP binding, a positive is an observed enzyme–ligand association. Negatives are sampled enzyme–ligand pairs without an observed association in the source, balanced to the number of positives; they are assumed negatives, rather than experimentally established non-binders. For residue annotation, a positive position is annotated as an active or binding site, and an unannotated position is assigned zero. Incomplete annotation can therefore affect the interpretation of both tasks. EC labels additionally distinguish known positives from masked, unresolved descendants (Section 2.2).

The 36 case-study datasets contain 106–15,840 rows each. Of these, 24 are variant libraries of one enzyme scaffold, including the Protein Engineering Tournament [5], deep mutational scans, combinatorial libraries, and PafA kinetic measurements. The median Hamming distance to the within-dataset consensus reference is 1 substitution in 21 libraries, 2 in the galactose oxidase library, and 3 in the two three-site libraries (Appendix Table 3). These are separate medians, rather than a median expressed as a range, and the consensus reference need not be the experimentally defined parent. The other 12 datasets are panels of 12–146 distinct enzymes measured against shared substrates or assay conditions. Several datasets come from the same experimental study or report different endpoints for the same variants, so 36 datasets do not represent 36 independent campaigns.

### 2.2 Data Processing & Modeling

Each dataset is presented in a tabular format where each row contains a labeled target. Sequences are de-duplicated into one pool of 319,791 unique sequences shared by all tasks and are never truncated. ESM3 ran out of GPU memory on 620 of the 658 sequences with 2,645 residues or more, so they were removed and all models see the same 319,171 sequences; the constraint is the memory of a single extraction rather than a fixed length cutoff, and the other 38 long sequences, up to 2,860 residues, remain in the pool and set the maxima of Table 1 (Section 4). UniProt flags an entry as a fragment when the sequence covers only part of the protein. In the six tasks whose label describes the entire, unfragmented protein (EC number, *k*_cat_, *K*_M_, pH_opt_, binding, and solubility), we remove every row whose sequence occurs only under fragment accessions (5,443 rows under the random split, 3,357 of them from EC number, highlighted further in Appendix H). The residue tasks retain the fragments, since their labels are positions on the deposited sequence.

Several tasks have multiple measurements for the same enzyme. CatPred-DB keeps one value per enzyme–substrate pair, the maximum *k*_cat_ and the geometric mean of *K*_M_ over the reported measurements, which enters the model as a feature (Section 2.3). 57% and 65% of the enzymes have more than one row. Binding pairs each protein with a median of 2 and up to 14 ligands. The optimal pH of a row is the median of the values BRENDA reports for one entry, defined by its EC number, organism, and UniProt accession, so 17.0% of the rows combine more than one measurement. Rows with the same sequence and the same values are merged, and 313 sequences keep more than one row because different BRENDA entries report different values, 221 of them under different EC numbers. In Δ*T*_m_ and ΔΔ*G*, repeated measurements of one mutant stay as separate rows and only identical values are merged: 849 of the 2,312 Δ*T*_m_ mutants and 714 of the 2,518 ΔΔ*G* mutants have more than one row. Under both split types, all rows of one sequence fall in the same split.

Regression targets are standardized with the mean and scale of the training split, and most activities and kinetic constants of the case studies are first converted to log_10_ values (Appendix H). The binding positives are enzyme–ligand pairs observed in BioLiP, and each protein receives as many negatives, drawn at random from the ligands it is not observed to bind, so the task is balanced (Appendix I).

EC prediction must account for multifunctional enzymes and partial annotations: a single enzyme can catalyze multiple reactions, and a database may resolve only the upper levels of an EC number. For example, across the 279,501 Swiss-Prot entries we downloaded (304,416 total EC annotations), 7% of enzymes carry multiple EC numbers and 16% of annotations are incomplete, with unknown levels written as a dash (for example 3.2.2.-, three-quarters of which lack only the fourth level). We encode the EC numbers of a protein as a hierarchical multi-label target with one group of classes per level, so that the label of a protein is a binary vector over the EC classes seen in training: a class is positive (target 1) when the row annotates the protein with it, and is assigned target 0 otherwise unless the class is masked as unresolved. A zero is a database-label convention, not experimental evidence that the enzyme cannot catalyze that reaction. Each EC number of a protein is therefore a positive at every known level, for example 3.2.2.6 is a positive for the classes 3, 3.2, 3.2.2, and 3.2.2.6. An incomplete EC number is a positive down to its last known level, and the classes below it are masked from the loss and from the metrics rather than counted as negatives, so a protein annotated 3.2.2.- is never scored as a negative for 3.2.2.6. Entries with the same sequence and the same EC numbers collapse into one row, and sequences annotated with different EC numbers under different accessions keep one row per annotation.

*Random splits* assign each protein, including all its rows, to training, validation, and test with an 80%/10%/10% split. All measurements for one enzyme stay together in one split. In the mutation tasks, each mutant sequence is assigned independently, so variants of the same wild type can fall in different splits. *Similarity splits* cluster sequences with MMseqs2 [63] at 30% identity and 80% coverage and assign each cluster, including all its sequences, to a single split. This removes close homologs from the test set but not every distant one, because each sequence joins a single cluster representative. At most, 0.1% of proteins in the similarity-split test set have a hit at 90% identity or more, compared to 44.1% of the proteins in the random split, and 7.9–54.6% have one at 30% identity or more (Appendix H). The case-study datasets use one random 80%/10%/10% split by row, stratified on ten bins of the target. In 14 case studies, the same enzyme is measured on several substrates or conditions, and because the split assigns rows, 97 to 100% of their test rows have their sequence in training (Section 3.5).

### 2.3 Embeddings and Predictors

We evaluated 14 frozen PLMs ranging from 274M to 3.5B parameters, with embedding dimensions ranging 768–2560 (Appendix Table 12): Ankh and Ankh3 [2, 20], ESM2 [48], ESM3 [30], ESM-C [14], ProstT5 [32], ProtT5 [21], and Profluent E1 [36]. Per-residue representations come from the final hidden layer (of the encoder, for encoder–decoder models). As a reference without a PLM, the same heads are trained on 36 sequence descriptors computed with Biopython’s ProtParam [17, 26] and, for the residue tasks and attention pooling, on a one-hot encoding of each residue (Appendix H); we call it no-plm.

Global tasks use the pooled *D*-dimensional embedding. Tasks with a substrate, binding, *k*_cat_, *K*_M_, and 13 case studies append the 200 RDKit [44] 2D descriptors of the substrate, normalized to [0, 1] with DescriptaStorus [38]. Case studies with assay conditions append them as features: temperature, pH, buffer in PETase, and substrate concentration in the copper radical oxidase panel. The linear and MLP heads on mean-pooled inputs standardize each input column with the training mean and scale, except the substrate descriptors. Mutation tasks use the difference between the mutant and wild-type embeddings, which have the same length in every row: mean pooling subtracts the pooled vectors, and attention pooling pools the per-residue difference. Residue tasks use the full *L × D* matrix, with one binary label per position.

The neural heads predict EC numbers with one output group per level and a binary cross-entropy that skips masked classes, averaged within each level and then over the four levels. XGBoost predicts the four levels concatenated into one vector of 5,729 classes under the random split and 5,907 under the similarity split. It fits one multi-output tree model per block of 256 classes, on the proteins with no masked class in that block, and predicts a class that is constant in training as that constant.

*Mean pooling* averages over residues converting the *L D* per-residue matrix into a *D*-dimensional vector. *Attention pooling* adds a learned attention update to that mean: the mean of the layer-normalized residues attends over the residues and the result is added back, so a silent attention branch returns mean pooling exactly. For the residue tasks, one self-attention layer is applied over the residues. The *linear* head is one linear layer, while the *MLP* adds a hidden layer of 2,048 units (256 on residue tasks) with dropout 0.1; both are trained with AdamW (Appendix B). *XGBoost* [15] fits gradient-boosted trees on the mean-pooled vector, or on each residue’s vector. Mean pooling with the three heads and attention pooling with the two neural heads give five architectures, and one embedding with one architecture is a competitor. The 14 PLMs and the no-plm reference give 75 competitors, 150 runs per property task and 75 per case studies dataset, 4,200 runs in total. The MLP configuration was chosen once via search on *K*_M_ with esmc_300m (Appendix C) and used everywhere; therefore performance differences come from representations and not individual model tuning.

### 2.4 Metrics and Their Interpretation

Regression reports MAE, RMSE, *R*^2^, and Kendall’s *τ*. Classification reports accuracy, precision, recall, F1, AUROC, and average precision, denoted AUPRC throughout this paper. The primary per-task displays use Kendall’s *τ* for regression and AUPRC for classification. MAE and RMSE are evaluated on targets standardized using the training mean and scale, so they are expressed in training standard deviations, rather than the original biochemical units. Converting an error back to physical units requires the stored scale and, for log targets, explicit treatment of the logarithmic transform.

EC and residue classification metrics pool the unmasked label positions, rather than averaging equally over classes or proteins. They can therefore emphasize frequent EC labels or longer sequences. Precision, recall, and F1 use the fixed decision threshold of 0.5, while AUROC and AUPRC use probability scores. Unannotated sites and sampled binding controls are treated as negatives for evaluation, with the annotation caveats in Section 2.1. Appendix I specifies the score-histogram approximation and bootstrap computation.

### 2.5 Global Rating with Training-Noise Ties

Metrics on different task types do not share a scale, so we aggregate pairwise comparisons rather than raw scores. For each task and split, 1,000 paired bootstrap resamples compare competitors on the same sampled test observations. Regression uses Kendall’s *τ* (weight 0.75) and MAE (0.25); classification uses AUROC, AUPRC, F1, precision, and recall (0.2 each). A competitor wins only when the metric gap exceeds 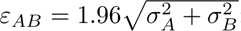; otherwise each competitor receives half a win. The training-noise scale σ is shared across PLMs with the same task, split, metric, head, and attention setting. It is estimated from a subset retrained with seeds 43 and 44, in addition to the published seed 42, using a median of seed standard deviations corrected for the small number of seeds (Eq. 1; 3,844 additional runs). Under independent normal training noise with known scales, the multiplier 1.96 corresponds to a 95% reference range for a score difference. Empirical seed gaps have heavier tails, so this is a calibrated comparison margin rather than a guaranteed 5% error rate (Appendix E).

We fit weighted Bradley–Terry ratings to these fractional win outcomes [12, 50]. The implementation uses logistic pairwise fitting with half-credit for ties, rather than the separate tie-probability likelihood of Rao–Kupper [56]. Fits to individual bootstrap rounds provide 95% percentile intervals. These intervals reflect test-observation resampling conditional on the published fitted models, task set, and weights; training variability influences the tie margin but is not itself resampled into the intervals. Our property weights prioritize reaction and catalytic-site annotation, with a 40%/60% random/similarity split within each task (Table 6). Case-study datasets have equal weights and a separately fitted leaderboard. Equal task weights provide a sensitivity analysis, rather than a unique alternative definition of representation quality.

### 2.6 Leaderboard, Cards, and Dataset Ingestion

Benchmarks fall out of date as new experimental campaigns are published, while manual curation and full re-embedding are costly. Enzymado uses structured provenance cards and an append-only ingestion workflow so new datasets can be added without re-embedding existing sequences (Appendix M).

Every component of Enzymado is described by a card. The benchmark card holds the task registry, split files, filters, and the rating configuration (metric, task, and split weights and the training-noise thresholds *ε_AB_* of Section 2.5). The rating code reads its settings from this card, and each leaderboard file stores the values it used. Each of the 46 dataset cards has two parts. The first is computed from the processed files: rows, distinct proteins, sequence lengths, label distributions per split, missing and duplicated values, and sequence redundancy from MMseqs2 clustering at 50% identity. The second records what cannot be computed: the assay and its conditions, the replicate design, the label noise reported by the source, and a reconciliation of every row removed or corrected between the source and our files. Each of the 14 PLMs has a card with its architecture, pretraining data and objective, parameter count, embedding dimension, and the exact way we extract its embeddings. This includes the special tokens we strip and the inputs we omit, such as E1’s retrieved homologs and Ankh3’s task prefix. The model cards contain no Enzymado results, so an explanation built from them does not simply restate the leaderboard.

Publication-derived card fields carry source quotations and page references, while stored provenance audits record checks against source text. Matching quoted text supports traceability; it does not establish that an interpretation is correct. The cards and converters were prepared with LLM assistance and author review. The ingestion process appends new sequences without renumbering existing ones, so existing embeddings can be reused and only added sequences require extraction. The agent assisted with adding 28 case-study datasets and identifying a mislabeled temperature row in the PET hydrolase panel [22] and colliding enantiomer SMILES in a galactose oxidase screen (Appendix M.4).

The leaderboard is hosted on Hugging Face and has two boards, properties and case studies. Each ranks the 75 competitors with their 95% interval and the range of ranks they took across bootstrap rounds. The two boards are fitted separately, so a rating on one is not comparable with a rating on the other. A results tab shows the primary metric of every competitor on every dataset, filtered by split, head, and metric. An AI assistant on the website answers questions about the results and takes every number from tools that retrieve the published results.

## 3 Results and Discussion

### 3.1 How Well Do PLM Representations Perform on Individual Enzyme Tasks?

PLM representations excel at classifying what reaction an enzyme catalyzes and which residues catalyze it, but remain weak at ranking enzymes by quantitative properties, with no single PLM dominating across tasks (Fig. 3). Under the similarity split with an MLP head and mean pooling, the best PLM reaches an AUPRC of 0.95 on active-site residue prediction and 0.84 on EC number prediction, compared with 0.04 and 0.14 for the no-plm baseline. On the five regression tasks, however, Kendall’s *τ* stays below 0.50 for every PLM, reaching only 0.25 on *k*_cat_ and surpassing no-plm by just 0.01 on *K*_M_ (0.43 versus 0.42). Under random splits, every PLM scores between 0.976 and 0.997 AUPRC on active sites and EC number, indicating that improvement on functional tasks lies in generalizing to distant homologs, whereas kinetic tasks require better representations or cleaner assay data.

**Figure 3:**
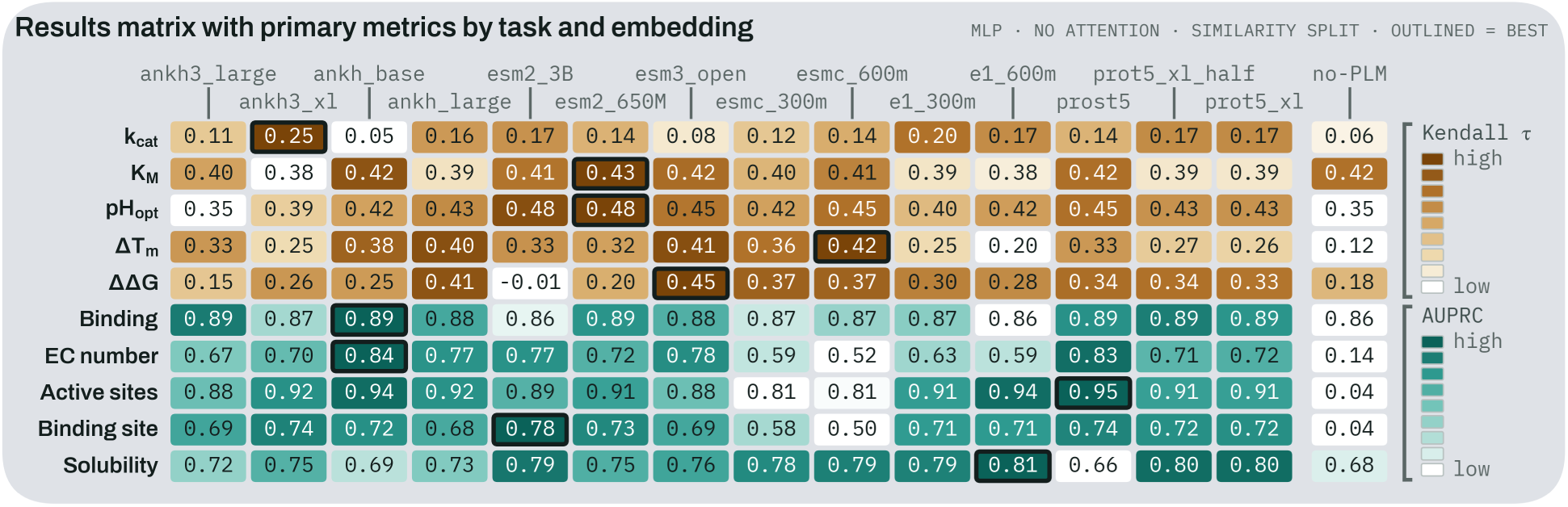
Results matrix with primary metrics by task and embedding. Primary metric per task and PLM (MLP head, mean pooling, similarity split): AUPRC for classification, Kendall’s *τ* for regression. Color is scaled within each row. e1_300m and e1_600m are the Profluent E1 models.

No single PLM is the best choice for every task. Eight different PLMs lead the ten tasks (Fig. 3), every task leader ranks 12th or lower on at least one other task, and the median rank of task leaders across the remaining tasks ranges from 4th (ProstT5) to 10th (Ankh3-XL). Within individual tasks, the leader is also rarely separable from competitors once test-set sampling and training-seed noise (*ε_AB_*) are accounted for: a median of 6 of the 13 other PLMs cannot be distinguished from the top model (Appendix F). On binding, for example, all 14 PLMs fall between 0.861 and 0.893 AUPRC, and 40 of the 91 pairs differ by less than *ε_AB_*. On ΔΔ*G*, the score range is the widest (*τ* from 0.013 to 0.449), but training-seed noise is similarly large, leaving 63 of 91 pairs tied inside *ε_AB_*, compared with 31 on EC number. A PLM should therefore be selected for the task at hand rather than from the global rating, and the stability tasks require either larger datasets or multi-seed averaging to separate models reliably.

### 3.2 Which Representation Leads Across All Enzyme Properties?

When aggregated across all property tasks, representations pretrained with 3D protein structure and decoded by a nonlinear MLP head lead the benchmark (Fig. 4). Under our tiered task weights, ProstT5 [32] with a mean-pooled MLP head ranks first at 1661 rating points (95% interval 1649–1674), taking first place in 91% of bootstrap rounds, with only five other competitors having overlapping confidence intervals. When all ten property tasks are weighted equally, the lead passes to ESM3, the only other PLM pretrained with structure (Kendall’s *τ* = 0.80 between the two rankings; Appendix D). ProstT5’s lead comes from tasks governed by catalytic and binding residues: it wins 74% of its matches on active sites, 76% on binding sites, and 61% on EC number while losing at most 6%, whereas it loses 48% of its matches on solubility. ProstT5 is initialized from ProtT5-XL and trained further to translate between amino acid sequences and the 3Di structural alphabet of Foldseek [39]. With the same MLP head, it rates 27 points above ProtT5-XL with non-overlapping confidence intervals, providing the closest comparison among the benchmarked models for isolating structural pretraining. All ten top competitors use the MLP head (eight without attention), while the bottom five are the no-plm baselines (Appendix H).

**Figure 4:**
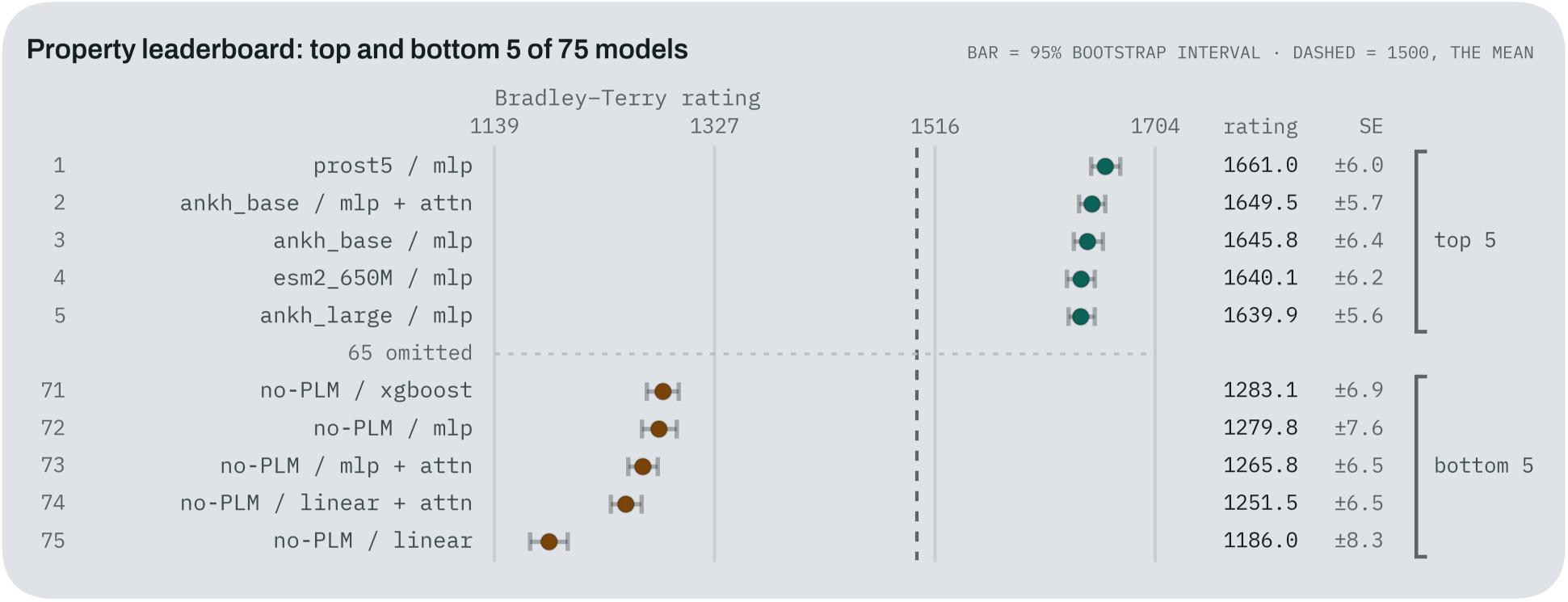
Enzyme properties leaderboard. The top and bottom five of the 75 competitors on the property leaderboard, with 95% bootstrap intervals; the bottom five are predictor heads of the no-plm baseline.

### 3.3 Are Newer and Larger PLMs Getting Better at Enzyme Tasks?

Neither recent release dates nor larger parameter counts yield better enzyme representations on the property benchmark (Fig. 5). Representing each PLM by its best predictor head, the six models released since December 2024 (ESM-C, Ankh3, and Profluent E1) hold the six lowest property ratings of the 14, and five have confidence intervals entirely below those of all eight older PLMs (Spearman *ρ* = 0.58 between release date and rating). This deficit is not an artifact of our task weights: taking the best architecture per PLM on each task, the newer models score fewer match points on eight of the ten tasks (losing most on pH_opt_, *K*_M_, EC number, and binding sites), tie on Δ*T*_m_, and lead only on solubility, where E1 ranks first under both splits. Under equal task weights, the six newer PLMs still occupy the six lowest ranks. Model size does not explain the gap either, as the newer PLMs span 274M to 3.48B parameters, covering the 453M to 2.84B range of the older models.

**Figure 5:**
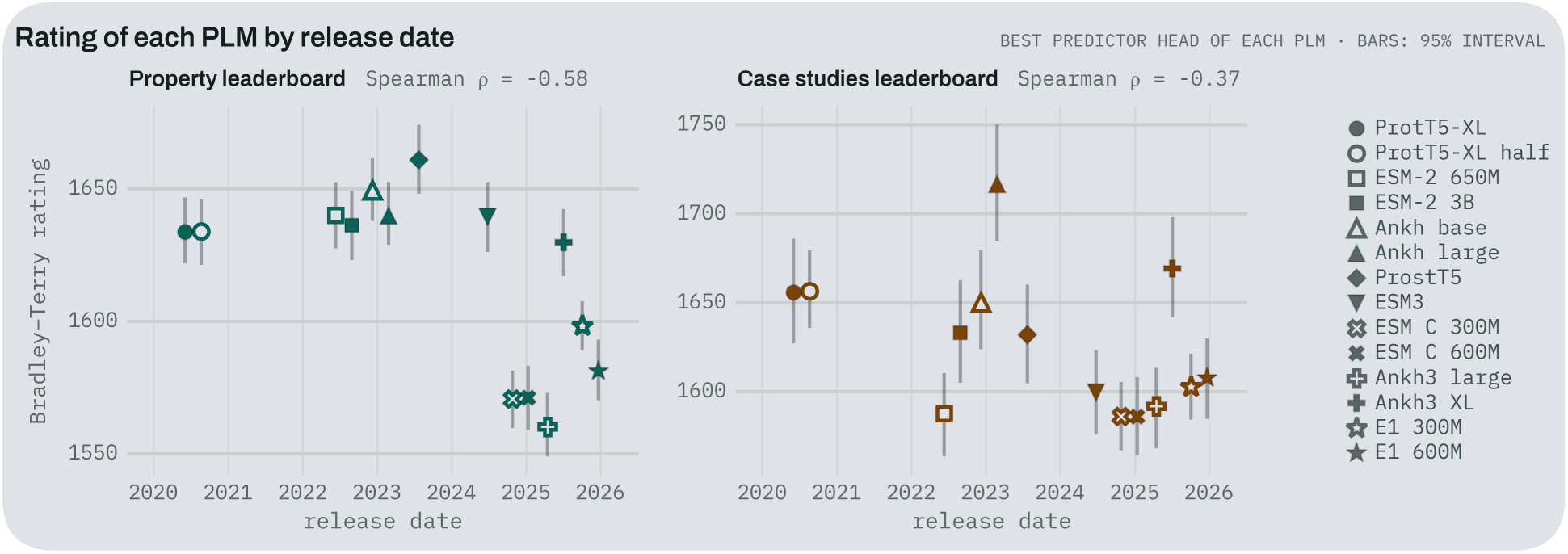
Rating of each PLM by first public release. Each PLM is represented by its best competitor, with its 95% interval; left, property leaderboard; right, case-study leaderboard. The six PLMs released since December 2024 hold the six lowest property ratings.

The two groups differ in pretraining: ESM-C adds metagenomic sequences to UniRef, E1 trains with retrieved homologs in context, and Ankh3 adds sequence completion to span denoising. None of these modifications improved frozen final-layer enzyme representations here, consistent with Li *et al.*, who showed that lower pretraining loss does not consistently transfer to downstream protein tasks [46]. Two of the newer families also run outside their pretraining setup (E1 without retrieved homologs and Ankh3 without its task prefix, though adding the prefix shifted probe scores by at most 0.014; Section 4), and whether they store more enzyme signal in intermediate layers remains untested. On the case-study leaderboard, the temporal trend keeps the same sign but is weaker (*ρ* = 0.37), with Ankh3-XL ranking second behind Ankh-Large. A newer release date is therefore no reason to switch PLMs for enzyme modeling without task-specific validation.

### 3.4 How Do Pretraining Objectives and Predictor Heads Shape Performance?

A representation’s rating depends as much on matching the predictor head to how information is encoded as it does on the PLM’s pretraining objective (Figs. 6 and 7). Grouped by training objective (Fig. 6), the two structure-trained models (ProstT5 and ESM3) reach the highest peak ratings when paired with an MLP head, while the encoder–decoder models trained with span denoising on sequences alone (ProtT5 and Ankh) maintain the highest median ratings across all five predictor architectures, ahead of encoder-only masked language models.

**Figure 6:**
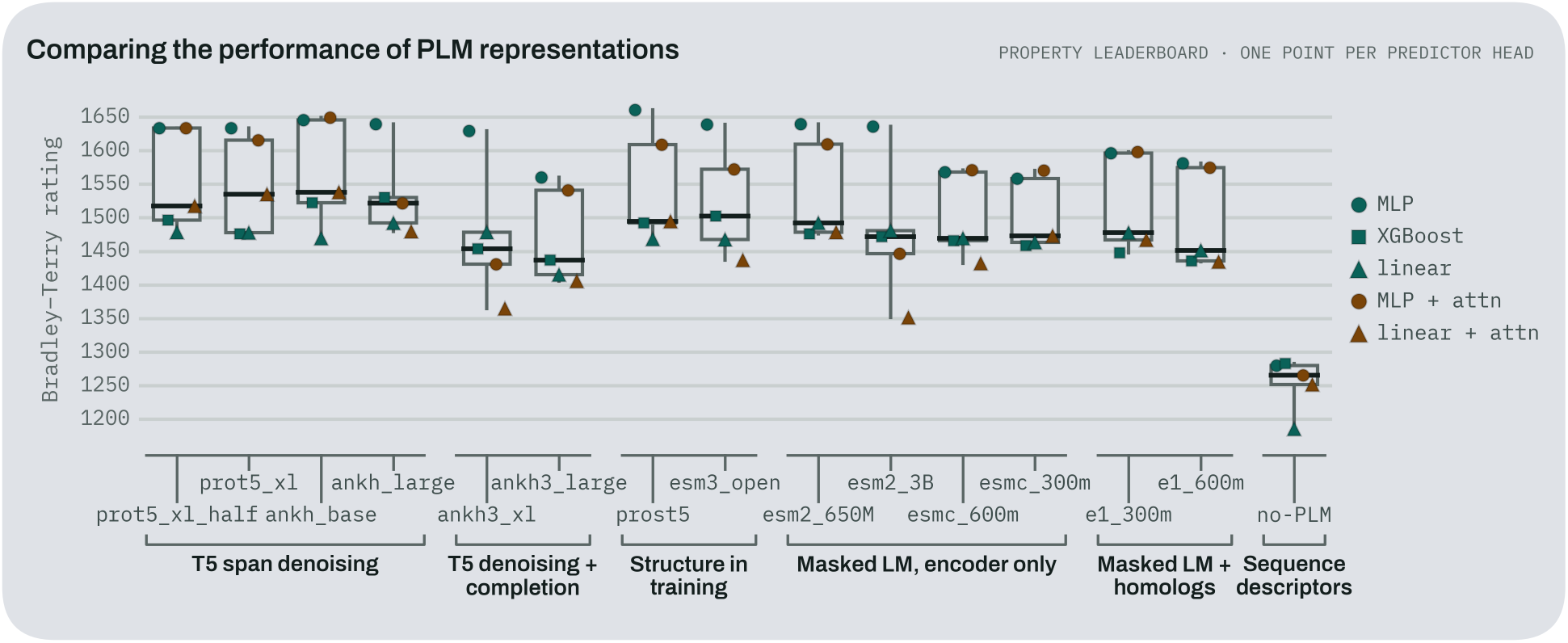
Rating of each PLM by architecture. Property leaderboard rating of the five architectures of each PLM, grouped by training objective. One point per architecture; boxes give the quartiles.

**Figure 7:**
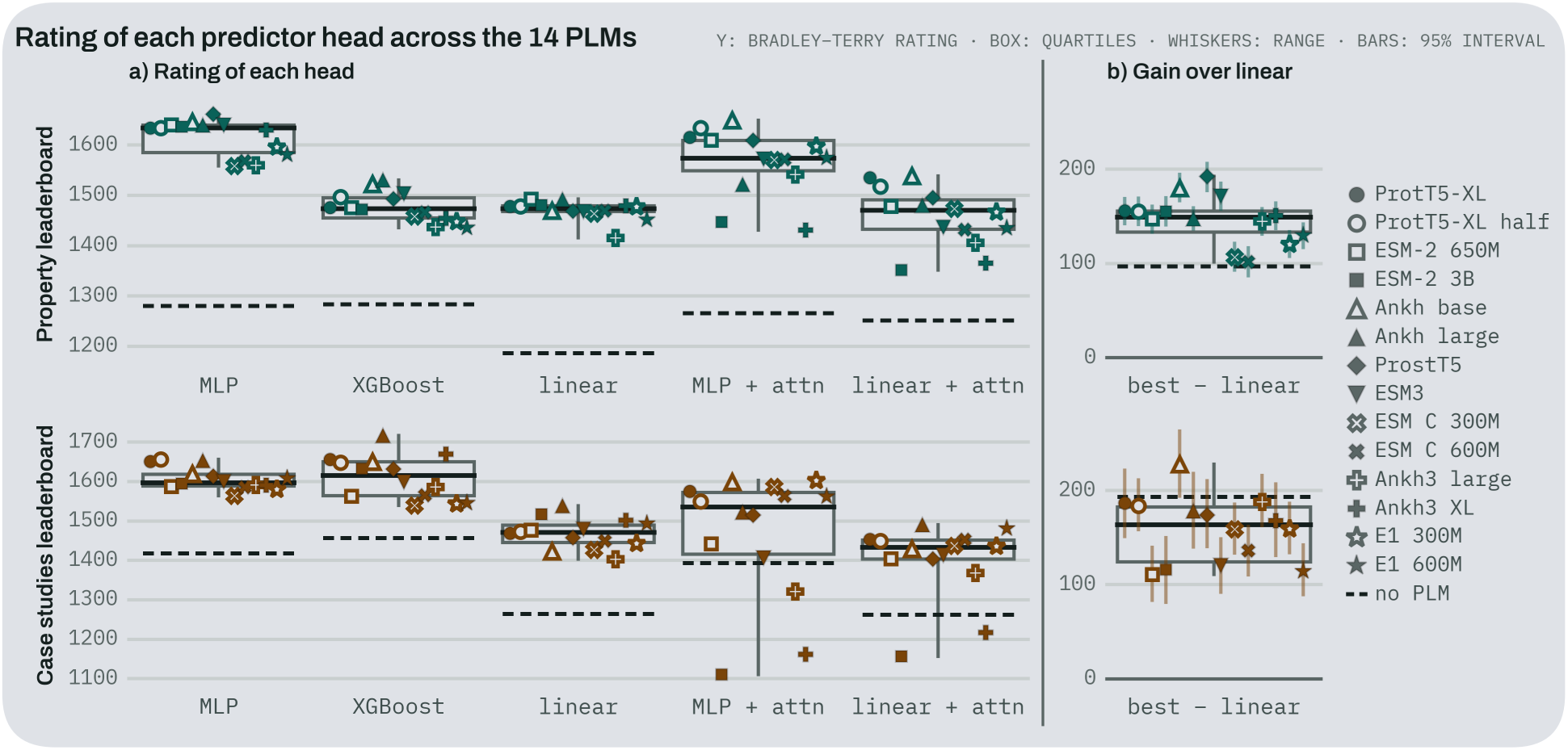
Rating of each architecture across the 14 PLMs. One point per PLM and the dashed line marks no-plm. Top, property leaderboard; bottom, case-study leaderboard.

Comparing linear and nonlinear heads tests how explicitly a representation encodes each property [10]. In the whole-protein scalar tasks, the mean-pooled linear and MLP heads stay within 0.04 of each other on every task except binding, showing that global enzyme properties are largely linearly accessible. On binding, the input concatenates the protein embedding and ligand descriptors, so a linear head scores each pair as an additive sum of protein and ligand terms and cannot capture protein–ligand specificity. In the residue tasks, however, the MLP reaches an average AUPRC of 0.95 in active sites and 0.83 in binding sites, compared with 0.67 and 0.50 for the linear head (Appendix G). Catalytic and binding residue identities are therefore present in the embeddings, but encoded nonlinearly (though part of the gap may also reflect that per-residue inputs are unstandardized and only the MLP applies dropout and weight decay). On the property leaderboard, every PLM gains 102 to 193 rating points with its best head over its linear probe (Fig. 7), and the two rankings agree only weakly (Spearman *ρ* = 0.46). Evaluating with linear probes alone reorders the leaderboard: ESM2 650M and Ankh-Large lead, while ProstT5 and ESM3 drop to 10th and 11th of 14 despite ranking first and fifth with an MLP. Because residue tasks carry high weight, predictor head selection shifts ratings as much as PLM selection. Finally, some tasks reward inputs beyond the protein representation: in *K*_M_ and binding, no-plm with substrate descriptors comes within 0.03 to 0.04 of the best PLM, and in residue tasks, homology-transferred Swiss-Prot annotations partly reward recognizing close relatives.

### 3.5 Do Database Benchmark Rankings Transfer to Protein Engineering Campaigns?

Rankings from the property leaderboard transfer only partially to experimental protein engineering campaigns (Spearman *ρ* = 0.42) because campaign datasets divide into two distinct modeling regimes. In 14 of the 36 case studies (12 multi-enzyme panels and two variant libraries assayed across multiple substrates), each sequence appears in multiple rows under row-stratified splits. A predictor can score well on these datasets simply by recognizing an enzyme seen in training and partitioning the substrate or condition features, which favors decision trees and requires no PLM: XGBoost beats the MLP on 13 of the 14 datasets, and the best no-plm competitor comes within a median of 0.01 in Kendall’s *τ* of the best PLM. In the remaining 22 case studies, which are single-enzyme variant libraries where every test sequence is an unseen mutant differing by a few substitutions, the representation must capture mutational effects: XGBoost beats the MLP on only one dataset, and the best PLM outperforms no-plm by a median of 0.21 *τ*. The 14 multi-row panels explain why XGBoost competitors take seven of the top ten spots on the case-study leaderboard, led by Ankh-Large at 1717 points (95% interval 1686–1749). As campaign test sets are small (10 to 1,664 rows), case-study rating intervals are also roughly twice as wide as on the property leaderboard. Until multi-substrate panels are split by enzyme, the 22 variant libraries are the subset that directly measures sequence representation quality.

### 3.6 What Mechanistic Hypotheses Do the Model and Dataset Cards Reveal?

Why one model beats another depends on both how the representation was trained and what the assay actually measures. Our model cards record architecture, pretraining data, objective, and extraction details, with zero Enzymado scores or rankings. That separation lets us pair a model attribute with a dataset attribute to form outcome-blind, testable hypotheses across all 56 task-splits (10 property tasks across two splits, plus 36 case studies) before looking at the leaderboard (Appendix M.5):

#### H1: Intra-family scaling and substrate feature dilution

Global parameter count did not predict overall ratings, yet comparing smaller and larger checkpoints within the same PLM family (ESM2, Ankh, Ankh3, ESM-C, and Profluent E1) shows a clear split depending on whether a substrate is in the input. Of the 56 task-splits, 37 take only the protein sequence and 19 append 200 RDKit substrate descriptors (Layout B). When protein width doubles while those 200 descriptors stay fixed, the substrate’s share of the input vector drops from 20.7% (200/968) to 7.2% (200/2,760). With a mean-pooled MLP head, the larger checkpoint beats the smaller one on 30 of 37 protein-only splits (81.1%, median gain +0.0091), then flips on Layout B, winning only 3 of 19 splits (15.8%, median change 0.0102). That reversal holds within every family and in the linear head (30/37 versus 4/19), while tree splits in XGBoost still benefit from scaling on protein-only tasks (31/37, 83.8%) and stay flat on Layout B (9/19, 47.4%, median 0.0008). A clean follow-up test is to project all protein embeddings to a shared width before concatenating substrate features.

#### H2: Embedding width and attention-pooling overfitting

Attention pooling learns query, key, value, and output projections over the *D*-dimensional residue vectors (`d_model`, *D* [768, 2,560]), so its parameter count grows as (*D*^2^): an attention block at *D* = 2,560 (ESM2 3B and Ankh3-XL) fits 11.1 times as many parameters per residue as one at *D* = 768 (Ankh-Base). Across all 14 PLMs, wider embeddings consistently lose more from adding attention on both the property (*ρ* = 0.91) and case-study (*ρ* = 0.86) leaderboards. At *D* = 768 (Ankh-Base), attention helps: it lifts the linear head by +69.0 rating points and the MLP head by +3.7 points (second overall on properties). At *D* = 2,560, adding attention to the MLP costs 189.6 and 198.6 points on properties and 484.1 and 428.9 points on case studies, dropping those models into the bottom four PLM spots on both boards. Down-projecting to a fixed attention bottleneck (or adding stronger regularization) can test how much of this drop is simple parameter overfitting.

#### H3: Structural pretraining and expression decoupling

ProstT5 starts from ProtT5-XL (keeping the same 24-layer, *D* = 1,024 architecture) and trains further to translate between amino acid sequences and Foldseek 3Di structural tokens, giving us our closest pair for isolating structural pretraining. Using the dataset card field is_expression_decoupled, we split the benchmark into 37 purified or concentration-normalized task-splits and 19 splits where the readout also depends on cellular abundance or expression (DeepSol and split-GFP solubility, VAMP-seq abundance, unnormalized lysate screens, and growth-coupled scans). With the mean-pooled MLP, ProstT5’s biggest gains over ProtT5-XL all land on expression-decoupled tasks: EC number (+0.103 AUPRC on the similarity split), Δ*T*_m_ (+0.063 *τ*), pH_opt_ (+0.040 *τ*), and active sites (+0.039 AUPRC). Its biggest losses land on expression-confounded tasks: DeepSol solubility (0.138 AUPRC on similarity), amidase growth selections (0.065 and 0.054 *τ*), and split-GFP solubility (0.055 *τ*), moving ProstT5’s mean rank among the 14 PLMs from 7.74 on expression-confounded tasks to 5.64 on expression-decoupled ones. One caveat remains: a ProtT5-XL checkpoint given equal extra sequence-only training would be needed to separate the 3Di objective from the extra compute.

#### H4: Protein multiplicity and sequence memorization

From the computed card counts, protein multiplicity *M* = rows/distinct_proteins measures how many times each enzyme repeats across substrates or conditions: *M* = 1.0 when every row is a unique sequence (22 single-enzyme variant libraries and 3 property datasets), rises just above it where a sequence keeps more than one entry or measurement (1.0002 in EC number, 1.03 in pH_opt_, 1.44 in Δ*T*_m_, and 1.49 in ΔΔ*G*), and reaches multi-substrate or multi-condition panels (up to 30.0 in PETase, 34.8 in galactose oxidase, and 96.0 in the esterase panel). Across all 56 task-splits, higher *M* tracks XGBoost’s lead over the MLP (*ρ* = +0.73, and +0.74 within case studies) and erodes the best PLM’s lead over no-plm (*ρ* = 0.73, and 0.79 within case studies). Splitting the 36 case studies by *M* also clarifies why the two leaderboards disagree: on the 22 variant libraries (*M* = 1.0), where every test row is an unseen mutant, T5 span-denoising models take the top four mean ranks (Ankh-Large at 4.61, Ankh-Base at 4.80, ProtT5-XL half at 4.95, and ProtT5-XL at 5.02, ahead of ProstT5 at 7.07, ESM2 3B at 8.07, and ESM3 at 8.73). On the 14 panels (*M >* 1.0), ProstT5 (6.04), ESM2 3B (6.57), and ESM3 (6.71) move to the front and XGBoost beats the MLP on 13 of 14 datasets.

#### H5: Assay noise and STRENDA completeness on kinetic tasks

Comparing noise_regime and strenda_ conditions across kinetic tasks helps explain why PLMs barely beat simple descriptors on database kinetics. CatPred-DB pools BRENDA and SABIO-RK measurements from many labs with 0 of 7 STRENDA conditions controlled or given to the model (an estimated inter-lab MAE floor of 0.38 to 0.71 log_10_ units), while the microfluidic PafA datasets hold all 7 of 7 STRENDA conditions fixed on one platform (10 to 15 replicates per variant, 95% CI 0.07 log units). When assay conditions are held fixed, the gap between the best PLM and no-plm opens up: on *k*_cat_, the PLM lead grows 5.0-fold from +0.031 *τ* on CatPred-DB (random split) to +0.155 *τ* on PafA, and reaches +0.266 *τ* (8.5-fold) on *k*_cat_/*K*_M_ and +0.077 *τ* (2.8-fold) on *K*_M_. Across all 46 regression task-splits, single-lab controlled assays (*n* = 38) also reach a higher median best-PLM *τ* than inter-lab aggregated datasets (*n* = 8, 0.543 versus 0.471), though protein diversity and split structure still differ between the two groups.

Taken together, *H1*–*H5* show why we pair leaderboard ratings with outcome-blind cards: a single score tells you which model won, while crossing model choices (`d_model`, pretraining_objectives) with assay and split structure (has_ligand_smiles, is_expression_decoupled, multiplicity *M*, and STRENDA completeness) helps separate what the representation actually learned from feature dilution, head overfitting, sequence memorization, and experimental noise (Appendix M.5).

## 4 Limitations

Enzymado is designed around how protein engineering labs use PLMs in practice: as plug-and-play feature extractors where frozen sequence embeddings are fed into lightweight predictors without per-model hyperparameter tuning or multi-GPU fine-tuning. That focus, together with the compute cost of exhaustive grid coverage (2,850 GPU-hours and 4.6 TB of stored embeddings), leaves fine-tuning, per-model hyperparameter sweeps, intermediate layers, start tokens, max pooling, alternative aggregation orderings [7], structure- or alignment-input PLMs, and contrastive [80] or ranking [69] losses untested, and it runs two PLMs outside their pretraining setup. Our substrate-conditioned tasks also concatenate 2D RDKit descriptors, which cannot distinguish stereoisomers, and stop short of full reaction prediction [1]. The property datasets inherit database biases, among them an oxidoreductase-heavy kinetic set and only 91 and 67 wild types behind Δ*T*_m_ and ΔΔ*G*, while our ingestion gate deferred over 50 candidate case studies. Two splitting caveats bound the conclusions: the 30% similarity split leaves distant homologs in the test set, and the 14 multi-substrate case studies are split by row, which rewards enzyme recognition over generalization (Section 3.5). Appendix J details each of these.

## 5 Conclusion

Enzymado shows that current PLM representations already capture what reaction an enzyme catalyzes and which residues catalyze it, while revealing that no single PLM dominates across all tasks. A representation should therefore be chosen for the task and assay regime at hand rather than by release date or parameter count, as the six PLMs released since December 2024 hold the six lowest property ratings. Averaged across the property suite, representations pretrained with 3D structure coupled to a nonlinear MLP head lead the benchmark (ProstT5 under our tiered weights and ESM3 under equal weights), and the choice of predictor head shifts ratings as much as the choice of PLM because catalytic residue information is encoded nonlinearly.

As new PLMs introduce simultaneous changes to training corpora, architectures, and multi-component loss functions, it becomes difficult to isolate which innovations actually drive downstream performance. Pairing our noise-calibrated evaluation with outcome-blind model and dataset cards turns these confounding design choices into testable hypotheses and points to concrete directions for building better enzyme representations.

### Compact representations for low-data regimes

Protein engineering campaigns operate on hundreds to a few thousand variants and often condition on substrates, and there wider embeddings are counterproductive: going from *D* = 768 to *D* = 2,560 dilutes appended substrate descriptors in neural heads and causes (*D*^2^) parameter overfitting in attention pooling, so future PLMs should prioritize information-dense representations over wider hidden states.

### Multi-component objectives and assay-aware pretraining

Continual pretraining on 3D backbone tokens (ProstT5) improves purified catalytic and stability assays at the expense of expression-confounded readouts, whereas span denoising (Ankh and ProtT5) leads on single-enzyme variant libraries, so balancing sequence, structure, and alignments through multi-component, contrastive, or ranking objectives can test whether one representation captures both local mutational effects and active-site geometry without losing sequence composition signals.

### Joint enzyme–substrate and condition representations

Database kinetic benchmarks underestimate PLM utility because they aggregate measurements across unrecorded conditions, and fixing all seven STRENDA conditions on one microfluidic platform raises the PLM advantage over sequence descriptors five- to eight-fold, so representations that jointly encode enzyme, substrate, cofactors, and conditions offer a direct route to quantitative kinetic prediction. Encoding the enzyme, its substrate, and the assay conditions as one graph is a natural starting point [8, 28, 79], since learned graph representations already outperform the fixed descriptors used here on both molecules and proteins.

Covering every dataset and modeling choice is not possible in a static study, and global rankings inevitably reflect which tasks are weighted most heavily. To make these findings directly actionable, Enzymado exposes its leaderboards, per-split results, and machine-readable model and dataset cards as programmatic data entry points alongside agentic workflows. This allows both practitioners and AI coding agents to select an evidence-based representation and predictor head for a specific assay regime (such as a low-data single-enzyme mutational screen or a substrate-conditioned panel), continuously ingest new experimental campaigns, and systematically query the cards to test new hypotheses on what drives enzyme modeling performance.

## A Data Sources

The 10 property datasets come from six sources and hold 593,011 rows over 257,722 distinct sequences (Table 1). Five are regression and five classification tasks, and each is evaluated under both the random and the similarity split at 80/10/10 (Section 2.2). Eight datasets describe the whole protein, whereas the two residue datasets label each position. Their sizes differ by a factor of 70, from 3,338 rows in Δ*T*_m_ to 233,586 in EC number, and the three Swiss-Prot datasets contribute 81% of the rows. The table counts rows after the cleaning of Section 2.2, which includes removing the 620 sequences that ESM3 could not embed, and in Δ*T*_m_ and ΔΔ*G* its proteins are mutant sequences derived from 91 and 67 wild types.

## B Optimization Details

Table 4 lists the settings of each predictor head. The neural heads are trained with AdamW, using the same settings for every embedding, and the MLP settings come from the search in Appendix C. The MLP uses SiLU activations without layer normalization, the learning rate stays constant, the regression heads minimize the mean squared error, and attention pooling uses one attention head. The best epoch is chosen by early stopping: after every epoch, the predictions on the validation split are scored, and once 20 epochs pass without improvement, training stops and the best checkpoint is restored. On regression tasks this score is Kendall’s *τ*, while on binding, solubility, and EC number it is AUPRC. Both metrics depend only on the order of the predictions, so a single badly predicted outlier on a skewed target cannot decide the epoch. The residue tasks are the exception: with millions of validation residues, computing AUPRC every epoch is too costly in memory, so their early stopping monitors binary cross-entropy instead; the histogram used below for EC number could lift this limit in a later version.

**Table 4:** Head settings. LR: learning rate; WD: weight decay; patience in epochs, or in boosting rounds for XGBoost; subsampling gives the row and column fractions per tree. The MLP hidden layer has 256 units on the residue tasks.

| **Head** | **Tasks** | **LR** | **WD** | **Epochs** | **Pat.** | **Other** |
|---|---|---|---|---|---|---|
| linear | pooled scalar | 10<sup>−3</sup> | 0.1 | 200 | 20 | batch 256 |
| linear | EC number | 10<sup>−2</sup> | 0 | 200 | 20 | per-level outputs |
| linear | residue, mean | 10<sup>−3</sup> | 0 | 200 | 20 | loader batch 128 |
| linear | residue, attn. | 10<sup>−3</sup> | 0 | 200 | 20 |  |
| MLP | all | 10<sup>−3</sup> | 0.1 | 100 | 20 | 2,048 units, dropout 0.1 |
| XGBoost | scalar | 0.1 | – | 100 trees | 20 | depth 6 |
| XGBoost | EC number | 0.1 | – | 50 trees | 20 | depth 4, subsampling 0.8/0.1 |
| XGBoost | residue | 0.1 | – | 40 trees | 10 | depth 3, subsampling 0.5/0.25 |

EC number raises a similar problem, since its validation split holds more than 130 million label positions. Rather than keeping them all in memory, we accumulate the validation AUPRC in a 16,384-bin score histogram, which is the same estimator used for the reported AUPRC. Batching also differs under attention pooling: each loader batch holds 16 proteins and gradients are accumulated over 16 batches, which matches the effective batch size of 256 used with mean pooling, except on the residue tasks. XGBoost, in turn, applies early stopping on the same validation split, monitoring AUPRC on binary protein-level tasks, RMSE on regression tasks, and log-loss on EC number and the residue tasks.

The linear head has no hidden layer, dropout, or normalization, which makes it a floor for comparison: the gap between a representation’s linear and MLP scores shows how much that representation relies on a nonlinear head to separate a property. Attention pooling weights residues via learned relevance, which is a capability mean pooling lacks and one that could matter when a few catalytic positions determine a whole-protein property. Since the parameters are learned in every run, a head with attention pooling has more parameters than the same head with mean pooling, so the two poolings are compared at unequal capacity.

## C Hyperparameter Optimization for the MLP Head

Early in the project, we searched MLP settings on the *K*_M_ regression task with the esmc_300m embedding to find a configuration that trained reliably before running the full set of experiments. We used Latin hypercube sampling to draw 400 configurations from the space in Table 5. The MLP row of Table 4 is one of these configurations and is used for every representation and task, except that the hidden layer has 256 units on the residue tasks.

**Table 5:**
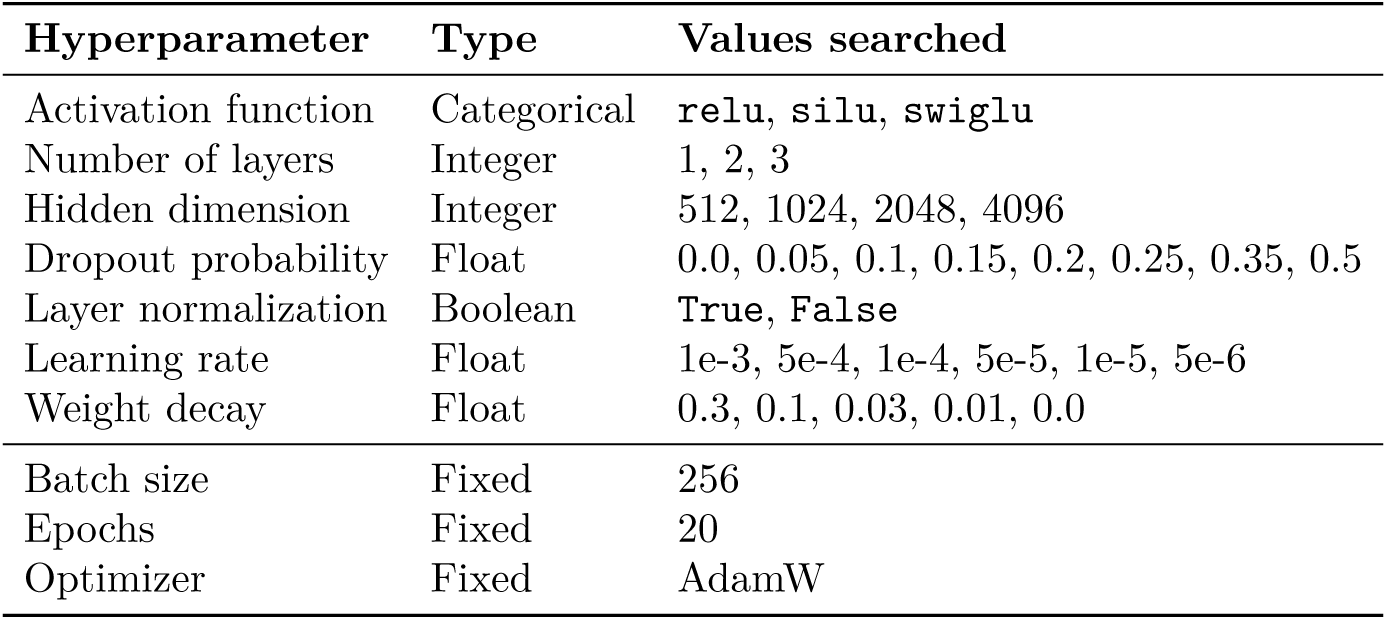
MLP hyperparameter search space, sampled on *K*_M_ with esmc_300m.

| **Hyperparameter** | **Type** | **Values searched** |
|---|---|---|
| Activation function | Categorical | relu, silu, swiglu |
| Number of layers | Integer | 1, 2, 3 |
| Hidden dimension | Integer | 512, 1024, 2048, 4096 |
| Dropout probability | Float | 0.0, 0.05, 0.1, 0.15, 0.2, 0.25, 0.35, 0.5 |
| Layer normalization | Boolean | True, False |
| Learning rate | Float | 1e-3, 5e-4, 1e-4, 5e-5, 1e-5, 5e-6 |
| Weight decay | Float | 0.3, 0.1, 0.03, 0.01, 0.0 |
| Batch size | Fixed | 256 |
| Epochs | Fixed | 20 |
| Optimizer | Fixed | AdamW |

## D Ranking and Confidence Intervals

### Resampling

For each task and split, we draw 1,000 bootstrap resamples of the test set and score every competing model on the *same* resamples, each indexed by a seed derived deterministically from the task and split. Before the resample indices are shared, the code verifies that each run’s ordered test sample identifiers match those of the first run, and any run whose test set differs is resampled independently. Pairing matters because two representations that differ by one point of AUPRC separate far more reliably when they share the noise of which test enzymes were drawn than when each is resampled separately. The reported intervals are the empirical 2.5th and 97.5th percentiles.

### Computing metrics under resampling

Recomputing every metric from scratch on each resample is infeasible for the largest tasks, where a single test split holds millions of annotated residues. We therefore compute classification metrics from per-sample confusion counts and score histograms, which can be resampled through reweighting, and regression metrics from the stored per-sample errors. Both approaches read the published prediction files, so each resample reweights real model outputs without rerunning any model.

### Matches and ties

Each resample yields one match per pair of models and rated metric. On regression tasks, Kendall’s *τ* has weight 3 and MAE has weight 1; on classification tasks, AUROC, AUPRC, F1, precision, and recall each have weight 1. RMSE, *R*^2^, and accuracy are reported but not rated. Within a resample, RMSE orders models exactly as *R*^2^ does, and MAE already captures the size of the error. Accuracy adds little because the all-negative classifier scores near one on rare labels, while on balanced labels precision, recall, and F1 already evaluate the same threshold. Weights within each task and split are normalized to sum to one. Each property task then carries its task weight (Section 2.5), with 40% from the random split and 60% from the similarity split, and each case-study dataset carries a weight of one from its single split. A match is won when the gap exceeds *ε_AB_*and tied otherwise; without training-noise estimates, this rule reduces to a strict comparison in which only identical scores tie. A second rule withholds a Kendall’s *τ* or AUROC cell when even the best model stays below 0.1 or 0.55, respectively, just above the no-skill values of 0 and 0.5; no cell met this condition in this version.

### Fitting

The ratings are the weighted Bradley–Terry maximum likelihood solution over the fractional match outcomes, centered on a mean rating of 1500. A fit pooled across all resamples gives the point ratings, and separate fits to each resample give the intervals. Because ties do not flip between resamples the way narrow wins do, the intervals are narrower with the noise threshold than without it (median half-width of 11.2 versus 19.7 rating points on the property leaderboard). These intervals therefore describe the uncertainty from the test set, not the joint uncertainty of test set and seed.

### Task weights

The property tasks are weighted by how directly they test knowledge of enzymes (Table 6). The first tier covers which reaction an enzyme catalyzes and which residues catalyze it; the second, how fast it turns over and where it binds; the third, substrate recognition under turnover and the ionization of catalytic groups; the fourth, folding and thermal stability, which all proteins share; and the fifth, the two tasks largely determined by the ligand descriptors or the expression system. Adjacent tiers are separated by an equal step, so no task outweighs another by more than a factor of five. The similarity split carries 60% of each task’s weight because predicting distant enzymes is the harder test. Weighting all tasks equally changes the order of the leaders, placing esm3_open first and ProstT5 fourth (Kendall’s *τ* = 0.80 against the official ranking); we keep the tiers because they reflect which tasks test enzyme function most directly.

**Table 6:** Task weights of the property leaderboard. *Share* is the fraction of the total weight; each task’s weight is split 40% to the random and 60% to the similarity split. The case-study datasets weigh equally.

| **Tier** | **Task** | **Weight** | **Share** | **What it tests** |
|---|---|---|---|---|
| 1 | EC number | 1.0 | 16.7% | which reaction the enzyme catalyzes |
|  | active sites | 1.0 | 16.7% | which residues catalyze it |
| 2 | *k*<sub>cat</sub> | 0.8 | 13.3% | how fast it turns over |
|  | binding sites | 0.8 | 13.3% | where it binds its ligands |
| 3 | *K*<sub>M</sub> | 0.6 | 10.0% | substrate recognition under turnover |
|  | pH<sub>opt</sub> | 0.6 | 10.0% | ionization of catalytic groups |
| 4 | ΔΔ*G* | 0.4 | 6.7% | folding stability, common to all proteins |
|  | Δ*T*<sub>m</sub> | 0.4 | 6.7% | thermal stability, common to all proteins |
| 5 | binding | 0.2 | 3.3% | largely decided by the ligand descriptors |
|  | solubility | 0.2 | 3.3% | largely decided by the expression system |

## E Measuring Model Performance Variability

The ratings account for two sources of noise: which test rows were drawn, handled by the paired bootstrap in Appendix D, and which seed trained the head, measured here. Training noise could be measured directly by repeating every run over several seeds, but a single pass over the grid took about 1,800 GPU-hours, and a single run on the residue tasks or on EC number with attention has a median duration of 2 to 8 hours (Appendix L). We therefore retrained a subset of runs with two additional seeds, for a total of 3,844 extra runs. This subset includes all 14 PLMs for the mean-pooled linear and MLP heads on the seven scalar property tasks and the 36 case-study datasets. For attention pooling, EC number, and the residue tasks, it includes four PLMs drawn at random from three groups of embedding dimensionality: one with up to 1,024 dimensions, two with 1,152 to 1,536, and one with 2,560. XGBoost without row or column subsampling is deterministic, so its *σ* is zero on the scalar tasks, as two extra seeds of one PLM on Δ*T*_m_ and ΔΔ*G* confirm; on EC number and the residue tasks, which subsample, *σ* comes from the same four PLMs.

For each combination of task, split, metric, head, and pooling, let *s_i_* be the standard deviation of PLM *i*’s scores over its *n_i_* seeds (*n_i_* = 3, or 2 when one run was left out). We define the training noise of that configuration as

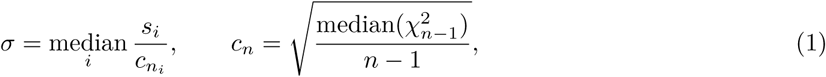

where the median is taken over the PLMs measured in that configuration and 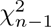 is the chi-squared distribution with *n* 1 degrees of freedom. Under normal noise, the standard deviation of *n* seeds has median *c_n_σ*, with *c_n_* = 0.83 for three seeds and 0.67 for two, so dividing by *c_n_* makes each term a median-unbiased estimate of *σ*. Taking the median over PLMs, a robust measure of scale [59], prevents one PLM with a collapsed seed from setting the tie threshold for an entire configuration. Regression runs with a training *R*^2^ below 1 are excluded from *σ* but remain on the leaderboard (221 runs, 90 of them on the *β*-glucosidase datasets). The resulting *σ* sets the tie threshold *ε_AB_* in Section 2.5.

The noise threshold uses one *σ* for all PLMs in a configuration. With only three seeds, each PLM’s standard deviation varies substantially by chance, even if all PLMs have the same underlying noise. To test whether one *σ* is sufficient, we simulated 14 PLMs with identical noise and compared the spread of their standard deviations with that of the real PLMs (Fig. 8, left). Across the 210 combinations of configuration and metric in which all 14 PLMs have three seeds, the real spread is close to the simulated one (median 0.62 versus 0.58), and 10% of the combinations exceed what chance explains, compared with 5% expected. One *σ* per configuration is therefore a fair simplification. Only `esm2_3B` is clearly noisier than the others, so its threshold is slightly too narrow. This check requires all 14 PLMs to have three seeds, which holds only for the mean-pooled heads on the scalar property tasks and case studies; for attention pooling, EC number, and the residue tasks, the shared *σ* is an assumption.

**Figure 8:**
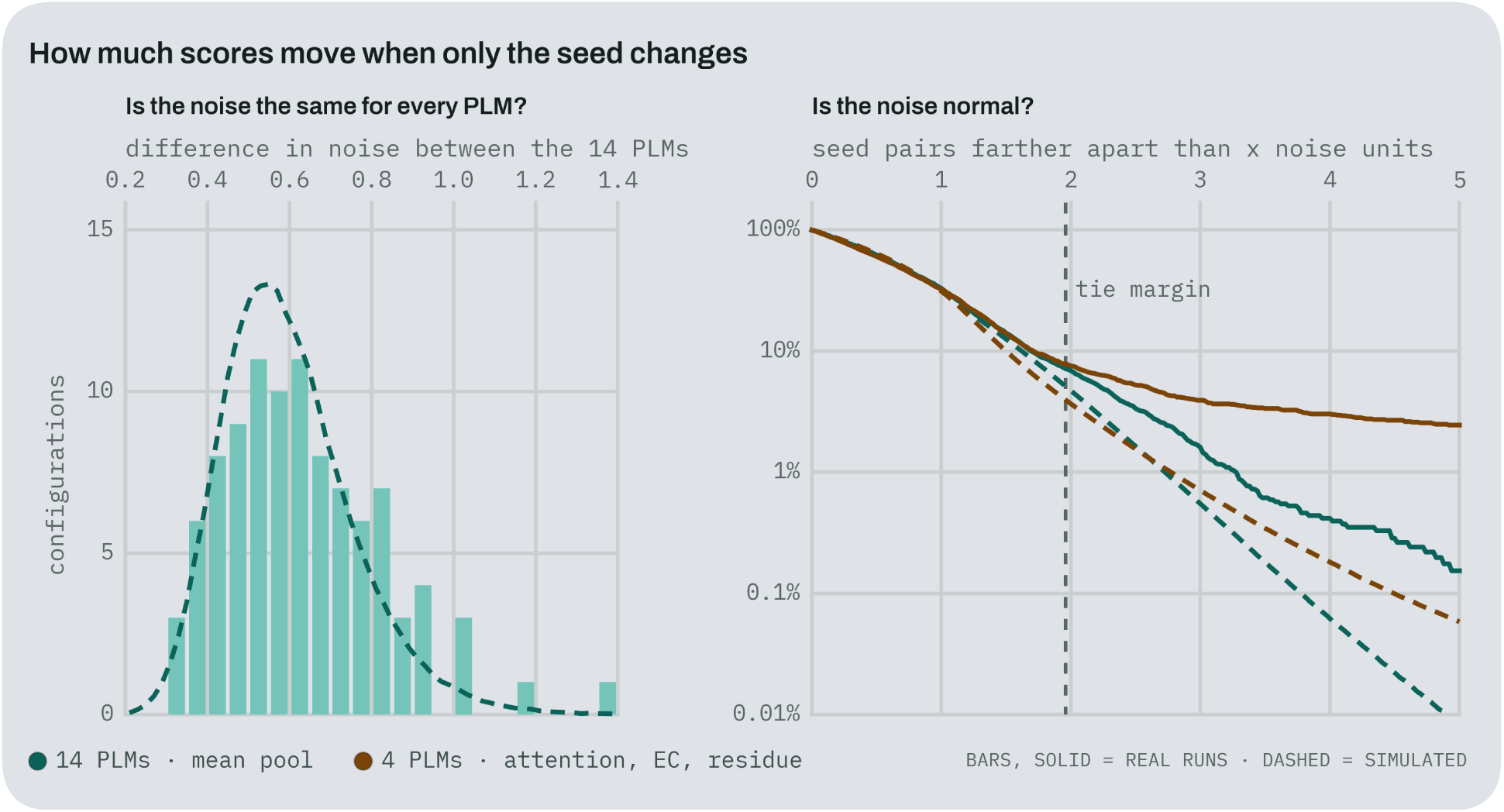
How much scores move when only the training seed changes. Left: the 210 combinations of task, split, head, and metric with mean pooling, the only setting where all 14 PLMs have three seeds. For each combination, the x-axis is the standard deviation, across the 14 PLMs, of the logarithm of each PLM’s seed standard deviation; 0 means all PLMs are equally noisy. Bars count the real combinations; the dashed curve is the count expected when the 14 PLMs share one noise level, from 5,000 simulations with normal noise and th<u>r</u>ee seeds. Right: every pair of seeds of the same PLM gives a gap between its two scores, measured in noise units of √2*σ*, the typical size of such a gap. The y-axis (log scale) is the share of pairs whose gap exceeds *x* units, and the vertical line is the noise threshold at 1.96. Solid lines are the real runs; dashed lines simulate normal noise with the same PLMs, the same seed counts, and *σ* estimated as in Eq. 1. Teal: the 14 PLMs with mean pooling; brown: the 4 drawn PLMs, with attention pooling or on EC number and the residue tasks.

The threshold also assumes normal noise, under which about 5% of seed pairs from the same model fall ou<u>t</u>side it. We tested this on the 13,280 seed pairs from the same PLM, measuring each gap in noise units of √2*σ* (Fig. 8, right). Small gaps match normal noise, but large gaps are more common than expected: 7.9% of pairs fall outside the threshold, compared with 4.8% in a simulation with normal noise and the same PLMs and seed counts. The excess comes from rare runs that land far from their other seeds, most of all in the configurations with four PLMs (8.7% against 4.0%), led by EC number with attention pooling. The threshold therefore covers about 92% of seed-to-seed gaps rather than 95%, and taking *σ* as a median prevents these rare runs from widening it. Finally, Table 7 shows how the share of ties, the ranking, and the rating range change when the multiplier 1.96 of the *ε_AB_* threshold is set to zero, halved, or doubled.

**Table 7:** Sensitivity of the ratings to the multiplier *z* of the noise threshold 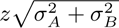; the ratings use *z* = 1.96, and *z* = 0 counts any difference as a win. *Ties*: share of all matches that end in a tie. *τ* : Kendall’s rank correlation between the ranking of the 75 competitors under that *z* and under *z* = 1.96. *Range*: rating of the first competitor minus rating of the last, in rating points.

|  | **Properties** |  |  | **Case studies** |  |  |
|---|---|---|---|---|---|---|
| *z* | **Ties** | ***τ*** | **Range** | **Ties** | ***τ*** | **Range** |
| 0 (strict) | 0.1% | 0.87 | 654 | 0.2% | 0.94 | 685 |
| 0.98 | 25.7% | 0.90 | 581 | 21.1% | 0.96 | 654 |
| 1.96 | 44.1% | 1.00 | 475 | 35.9% | 1.00 | 606 |
| 3.92 | 62.7% | 0.84 | 352 | 54.2% | 0.94 | 507 |

## F Paired Tests Against the Best PLM

The leaderboard ranks competitors across all tasks, whereas this appendix asks, for each task and split, how many PLMs cannot be distinguished from the best one. Using the baseline of Fig. 3 (MLP head, mean pooling), we compare the best PLM with each of the other 13 on the primary metric. Let *d_b_* be the score of the best PLM minus the score of the other PLM on bootstrap resample *b* = 1, 1000 from Appendix D. Because both models are scored on the same test rows in each resample, *d_b_* is a paired bootstrap of their gap. The other PLM is separable from the best when the 2.5th and 97.5th percentiles of *d_b_* exclude zero. Since each task and split involves 13 comparisons, we also compute the two-sided bootstrap *p*-value 2 min *P* (*d≤*0)*, P* (*d≥*0), where *P* (*d≤*0) is the share of resamples in which the best PLM does not lead, and adjust the 13 values with Holm’s procedure [34]. The last column of Table 8 adds training noise: the other PLM is separable only if the gap *d* on the full test set exceeds 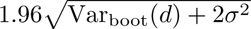. Here, Var_boot_(*d*) is the variance of *d_b_* across resamples, which captures test-set sampling noise, and 2*σ*^2^ is the noise that retraining adds to the gap between two models, each with the training noise *σ* from Eq. 1 for that task, split, metric, and baseline.

**Table 8:** Paired tests of the best PLM against the other 13, per property task and split (MLP head, mean pooling). *Best PLM* has the highest primary metric (Kendall’s *τ* for regression, AUPRC for classification) and *Score* is its test value. The last three columns count how many of the other 13 PLMs cannot be separated from it. *Test*: the 95% interval of the paired bootstrap differences contains zero, so only test-set sampling is considered. *Holm*: the two-sided bootstrap *p*-value, adjusted with Holm’s procedure over the 13 comparisons, is at least 0.05. *+Train*: the gap is within 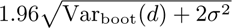, which adds the training noise *σ* of Eq. 1. A count of 0 means the best PLM is separable from every other, and 13 from none. These tests are reported only and do not enter the ratings.

| **Task** | **Split** | **Best PLM** | **Score** | **Test** | **Holm** | **+Train** |
|---|---|---|---|---|---|---|
| *k*<sub>cat</sub> | random | ankh3_large | 0.369 | 1 | 5 | 6 |
|  | similarity | ankh3_xl | 0.249 | 0 | 0 | 6 |
| *K*<sub>M</sub> | random | prost5 | 0.483 | 2 | 5 | 5 |
|  | similarity | esm2_650M | 0.428 | 1 | 4 | 11 |
| pH<sub>opt</sub> | random | ankh_base | 0.472 | 4 | 6 | 6 |
|  | similarity | esm2_650M | 0.483 | 1 | 1 | 8 |
| Δ*T*<sub>m</sub> | random | esm3_open | 0.470 | 10 | 13 | 11 |
|  | similarity | esmc_600m | 0.419 | 4 | 5 | 9 |
| ΔΔ*G* | random | ankh3_xl | 0.534 | 5 | 9 | 8 |
|  | similarity | esm3_open | 0.449 | 1 | 1 | 7 |
| binding | random | ankh_base | 0.923 | 1 | 2 | 5 |
|  | similarity | ankh_base | 0.893 | 6 | 8 | 7 |
| EC number | random | prot5_xl_uniref50 | 0.991 | 1 | 1 | 4 |
|  | similarity | ankh_base | 0.844 | 0 | 0 | 2 |
| active sites | random | esm2_3B | 0.997 | 0 | 0 | 0 |
|  | similarity | prost5 | 0.946 | 0 | 0 | 2 |
| binding sites | random | esm2_3B | 0.988 | 0 | 0 | 0 |
|  | similarity | esm2_3B | 0.778 | 0 | 0 | 0 |
| solubility | random | profluent_e1_300m | 0.868 | 7 | 8 | 9 |
|  | similarity | profluent_e1_600m | 0.806 | 7 | 10 | 8 |

Test-set sampling alone leaves a median of one PLM that cannot be separated from the best, or three after Holm’s correction. Adding training noise raises the median to six of 13. Only esm2_3B is separable from every other PLM under both sources of noise, on active sites under the random split and on binding sites under both splits. On Δ*T*_m_ under the random split, no PLM is separable from esm3_open after Holm’s correction.

## G Splits, Pooling, and Predictor Heads

The full grid allows for varying one factor at a time while holding the others fixed. Table 9 reports the split penalty for each task, averaged over every head, pooling, and embedding. The residue annotation tasks lose 0.14 to 0.16, consistent with many Swiss-Prot annotations having been propagated by homology. Table 10 restricts the pooling comparison to the linear and MLP heads, the only heads with both poolings, because averaging over all heads would compare two heads against three. Table 11 compares the heads under mean pooling.

**Table 9:** Split comparison. Primary metric averaged over all heads, poolings, and embeddings.

| **Task** | **Random** | **Similarity** | **Change** |
|---|---|---|---|
| *k*<sub>cat</sub> | 0.319 | 0.137 | −0.183 |
| *K*<sub>M</sub> | 0.459 | 0.390 | −0.069 |
| pH<sub>opt</sub> | 0.427 | 0.415 | −0.013 |
| Δ*T*<sub>m</sub> | 0.392 | 0.293 | −0.100 |
| ΔΔ*G* | 0.433 | 0.245 | −0.188 |
| binding | 0.839 | 0.821 | −0.018 |
| EC number | 0.901 | 0.588 | −0.313 |
| active sites | 0.759 | 0.615 | −0.144 |
| binding sites | 0.683 | 0.523 | −0.160 |
| solubility | 0.819 | 0.746 | −0.074 |

**Table 10:** Pooling comparison, restricted to the linear and MLP heads, averaged over embeddings and splits.

| **Task** | **Rows** | **Mean pool** | **Attention** | **Change** |
|---|---|---|---|---|
| EC number | 233,586 | 0.832 | 0.773 | −0.059 |
| binding sites | 157,124 | 0.662 | 0.680 | +0.018 |
| active sites | 89,329 | 0.809 | 0.748 | −0.061 |
| *K*<sub>M</sub> | 40,088 | 0.420 | 0.407 | −0.013 |
| binding | 28,316 | 0.824 | 0.801 | −0.023 |
| *k*<sub>cat</sub> | 22,693 | 0.232 | 0.213 | −0.019 |
| pH<sub>opt</sub> | 11,113 | 0.426 | 0.411 | −0.015 |
| ΔΔ*G* | 3,760 | 0.365 | 0.319 | −0.047 |
| solubility | 3,664 | 0.790 | 0.781 | −0.009 |
| Δ*T*<sub>m</sub> | 3,338 | 0.362 | 0.311 | −0.050 |

**Table 11:** Predictor head comparison under mean pooling, averaged over embeddings and splits. Best head per task in bold.

| **Task** | **Linear** | **MLP** | **XGBoost** |
|---|---|---|---|
| *k*<sub>cat</sub> | 0.220 | 0.243 | **0.252** |
| *K*<sub>M</sub> | 0.406 | 0.434 | **0.468** |
| pH<sub>opt</sub> | 0.419 | **0.433** | 0.430 |
| Δ*T*<sub>m</sub> | 0.351 | **0.372** | 0.367 |
| ΔΔ*G* | 0.348 | **0.382** | 0.328 |
| binding | 0.757 | 0.891 | **0.901** |
| EC number | 0.819 | **0.845** | 0.511 |
| active sites | 0.671 | **0.946** | 0.322 |
| binding sites | 0.499 | **0.825** | 0.332 |
| solubility | 0.785 | **0.795** | 0.772 |

## H Modeling Considerations

Table 12 lists the PLMs, and Table 13 gives the random split sizes. Proteins are assigned to splits in a seeded, deterministic order, with balancing on rows, so the 80/10/10 proportions hold for rows and only approximately for proteins. The kinetic datasets hold one value per assayed substrate and can therefore hold several per enzyme: *K*_M_ has 40,088 rows across 12,004 distinct proteins. The residue datasets have one row per protein, with a label for every residue.

**Table 12:**
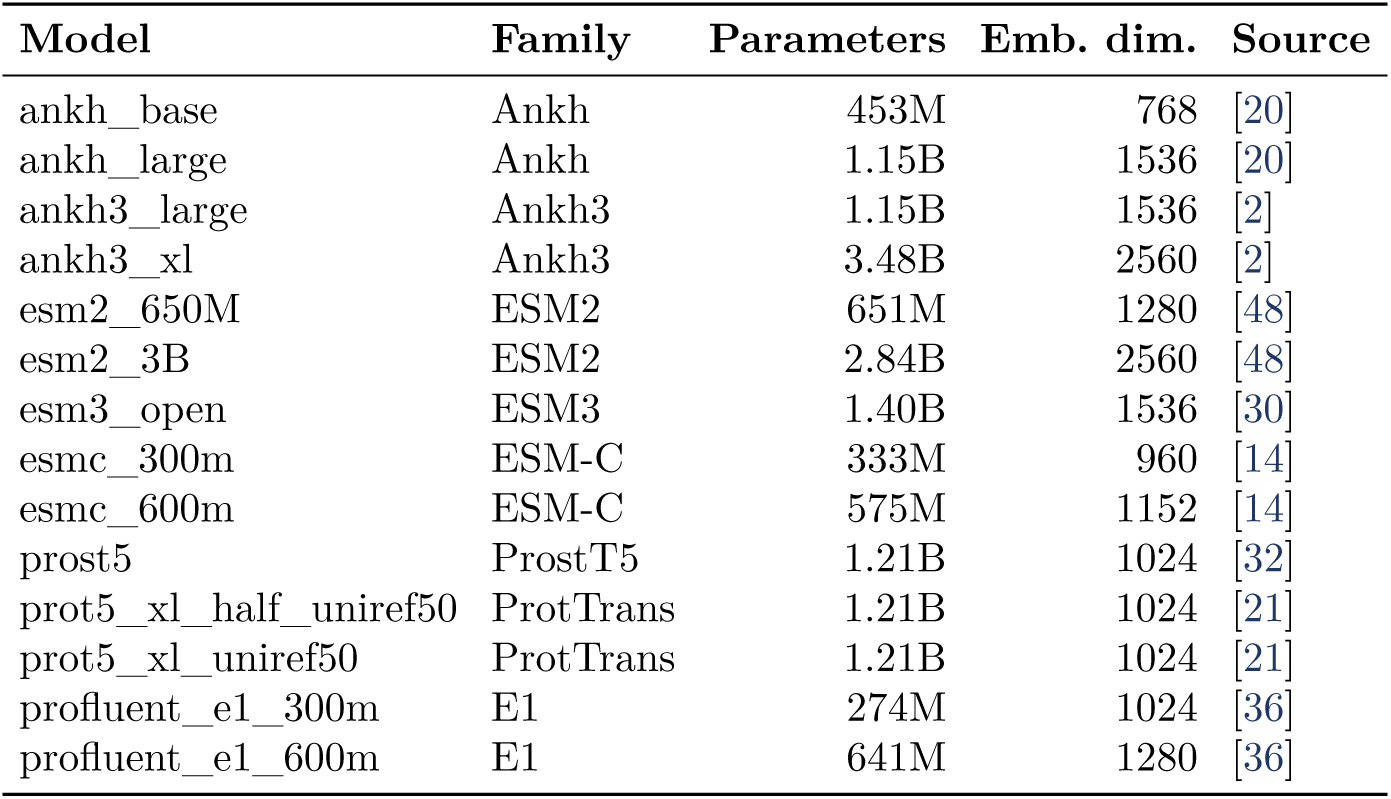
Protein language models. Parameters count only the encoder for the encoder–decoder models. prot5_xl_half_uniref50 is the encoder of prot5_xl_uniref50 stored in half precision.

| **Model** | **Family** | **Parameters** | **Emb. dim.** | **Source** |
|---|---|---|---|---|
| ankh_base | Ankh | 453M | 768 | [20] |
| ankh_large | Ankh | 1.15B | 1536 | [20] |
| ankh3_large | Ankh3 | 1.15B | 1536 | [2] |
| ankh3_xl | Ankh3 | 3.48B | 2560 | [2] |
| esm2_650M | ESM2 | 651M | 1280 | [48] |
| esm2_3B | ESM2 | 2.84B | 2560 | [48] |
| esm3_open | ESM3 | 1.40B | 1536 | [30] |
| esmc_300m | ESM-C | 333M | 960 | [14] |
| esmc_600m | ESM-C | 575M | 1152 | [14] |
| prost5 | ProstT5 | 1.21B | 1024 | [32] |
| prot5_xl_half_uniref50 | ProtTrans | 1.21B | 1024 | [21] |
| prot5_xl_uniref50 | ProtTrans | 1.21B | 1024 | [21] |
| profluent_e1_300m | E1 | 274M | 1024 | [36] |
| profluent_e1_600m | E1 | 641M | 1280 | [36] |

**Table 13:** Split sizes in rows under the random protocol, after the removal of UniProt fragments and the 620 sequences that could not be embedded.

| **Task** | **Train** | **Val** | **Test** |
|---|---|---|---|
| *k*<sub>cat</sub> | 18,163 | 2,257 | 2,273 |
| *K*<sub>M</sub> | 32,107 | 3,997 | 3,984 |
| pH<sub>opt</sub> | 8,887 | 1,118 | 1,108 |
| Δ*T*<sub>m</sub> | 2,670 | 335 | 333 |
| ΔΔ*G* | 3,010 | 375 | 375 |
| binding | 22,650 | 2,822 | 2,844 |
| EC number | 186,846 | 23,366 | 23,374 |
| active sites | 71,803 | 8,758 | 8,768 |
| binding sites | 126,252 | 15,452 | 15,420 |
| solubility | 2,932 | 365 | 367 |

### Log targets

Of the 36 case-study targets, 24 are modeled on a log scale. Four are already reported on a log scale by their source (the D-amino acid oxidase activity and expression scores, the aminotransferase specific activity, and the thiolase activity), and the other 20, activities, conversions, and kinetic constants, are converted to log_10_ values. Before the transform, an offset keeps zeros finite: 10*^−^*^3^ in ten datasets, 10*^−^*^4^ in the haloalkane dehalogenase panel, 1 in PETase, and, in the Protein Engineering Tournament datasets that contain zeros, the smallest positive value rounded to one significant figure. Galactose oxidase and the three PafA kinetic constants have no zeros and take no offset.

### Fragments

Fragment accessions were taken from UniProt release 2026_03 and account for 3,822 of the 248,802 accessions in the sequence index. Removing their rows did not move any remaining row to another split, and regression targets were restandardized on the remaining training rows.

### Homology between splits

We searched each test set of the eight non-mutation property tasks against its training set with MMseqs2 (*E <* 10*^−^*^3^, with at least 80% coverage of both sequences). Under the similarity split, at most 0.1% of test proteins have a training hit at 90% identity or more, and at most 9.3% have one at 50% or more; under the random split, these shares are 0.3–44.1% and 26.4–90.1%, respectively. At 30% identity or more, 7.9–54.6% of similarity-split test proteins and 60.5–96.6% of random-split test proteins have a hit. In the mutation tasks, no wild type spans two splits under the similarity split, whereas under the random split 98 and 100% of the Δ*T*_m_ and ΔΔ*G* test mutants share their wild type with a training mutant.

### The no-plm descriptors

The 36 pooled descriptors comprise amino acid composition, secondary structure fractions, molecular weight, isoelectric point, GRAVY, instability index, aromaticity, the two extinction coefficients, net charge at pH 7.4, and five statistics of the flexibility profile. For the residue tasks and attention pooling, each residue is represented by a 20-dimensional one-hot vector, whose mean over residues equals the amino acid composition.

### Class balance

Binding is 50.0% positive by construction because the BioLiP negatives were sampled to match the positives, and solubility is 56.1% positive. In the active-site dataset, 147,096 of 37.9 million residues are annotated, a positive rate of 0.39%; in the binding-site dataset, the rate is 2.51% of 67.7 million residues. A model that predicts no sites at all therefore scores above 99% accuracy on active sites.

### Accuracy under imbalance

In EC number, 0.07% of the scored label positions are positive, so predicting no class scores 0.9993. Across the 150 EC runs, accuracy spans 0.9967 to 1.0000, with three runs below that trivial score, while AUPRC spans 0.112 to 0.993 over the same runs.

### EC label structure

The EC vocabulary is fitted on the training split and passed to validation and test, so those splits never add a class unseen in training. The random protocol gives 7, 76, 271, and 5,375 classes at the four levels, and the similarity protocol gives 7, 76, 274, and 5,550, so the flat target of the tree head has 5,729 or 5,907 positions. An enzyme carries 1.09 EC numbers on average and at most 11, so about four positions of the flat vector are positive. This label shape, more than anything specific to enzymes, explains why EC number is the task with the largest similarity-split penalty and the protein-level task where gradient-boosted trees fall furthest behind the MLP (0.511 versus 0.845).

## I Labels and Metrics

### Binding labels

Positives come from the non-redundant BioLiP set. We keep entries with an EC number, a UniProt accession, and a ligand identifier, map each ligand identifier to a SMILES string through the wwPDB Chemical Component Dictionary, and remove polymer ligands (DNA, RNA, and peptides), single-atom ions, and SMILES that RDKit cannot parse. Protein–ligand pairs with the same sequence and SMILES are kept once. Each protein then receives as many negatives as it has positive ligands, drawn with a fixed seed from the dataset’s ligands that it is not observed to bind, which makes the task 50% positive. A rule that scores each test pair by the share of positives of its ligand in training reaches an AUPRC of 0.75 under the random split and 0.74 under the similarity split, against 0.50 by chance, so part of this task is decided by the ligand alone.

### Duplicate entries

Among the 279,501 Swiss-Prot entries with an EC number, 19,498 sequences appear under more than one accession. Entries with the same sequence and the same EC numbers are merged into one row, which removes 41,975 entries. The 52 sequences whose entries list different EC numbers keep one row per entry, all in the same split, so each row treats EC numbers found only in the other entry as negatives; these rows make up 0.04% of the dataset. In pH_opt_, rows with the same sequence and the same values are merged, and 313 sequences keep more than one row because different BRENDA entries report different values, 221 of them under different EC numbers.

### Metrics

All metrics are computed on the test split. Regression metrics use targets standardized with the training mean and scale, so MAE and RMSE are expressed in training standard deviations. Kendall’s *τ* is set to 0 when the predictions or labels are all equal. In classification, each scored position counts once: a row in binding and solubility, an unmasked class at any of the four levels in EC number, and a residue in the residue tasks. Precision, recall, F1, and accuracy are derived from confusion counts at a probability threshold of 0.5. AUROC and AUPRC are derived from histograms of the predicted probabilities in 16,384 bins: AUROC is the Mann–Whitney statistic, with a positive and a negative in the same bin counted as half a win, and AUPRC is the average precision accumulated from the highest bin down. Because the counts and histograms are stored per row, a bootstrap replicate (Appendix D) sums them over the resampled rows rather than recomputing predictions.

## J Limitations in Detail

Enzymado is designed around how protein engineering labs use PLMs in practice: as plug-and-play feature extractors where frozen sequence embeddings are fed into lightweight predictors without per-model hyperparameter tuning or multi-GPU fine-tuning. This practical focus, combined with the compute cost of exhaustive grid coverage (2,850 GPU-hours and 4.6 TB of stored embeddings), introduces several limitations that we plan to address in future versions.

### Modeling and optimization scope

We kept all 14 PLMs frozen, extracted only the final layer, pooled by the mean or a single attention block, and fixed one MLP configuration across all models and tasks. This leaves PLM fine-tuning, per-model hyperparameter sweeps (or Bayesian optimization to explore larger combinatorics efficiently), intermediate layers, start tokens, max pooling, and structure- or alignment-input PLMs untested. Alternative aggregation ordering also merits study; for example, applying an MLP per residue before pooling gives similar predictive accuracy while improving attribution interpretability [7]. Similarly, we trained with standard cross-entropy and mean squared error rather than contrastive learning [80], probabilistic calibration, or ranking losses [69]. Two PLMs also run outside their pretraining setup: E1 without retrieved homologs and Ankh3 without its task prefix ([NLU] or [S2S]), though adding the prefix changed linear-probe scores on 3,000 pH_opt_ and solubility proteins by at most 0.014.

### Reaction and dataset scope

Our substrate-conditioned tasks concatenate enzyme embeddings with 2D RDKit descriptors, which cannot distinguish stereoisomers, and we do not yet cover full reaction prediction or leakage-controlled multimodal reaction tasks as in Leak-CURBER [1]. The property datasets also inherit database biases: 28 to 29% of Swiss-Prot-mapped kinetic proteins are oxidoreductases and 4% are ligases (against 13% and 11% in Swiss-Prot), the Δ*T*_m_ and ΔΔ*G* sets span only 91 and 67 wild types, solubility keeps only 5% of DeepSol (3,664 enzymes), only 1.5% of active-site and 3.7% of binding-site labels cite experimental evidence, and excluding 620 sequences over 2,644 residues (under 0.5% of any task) omits giant polyketide and nonribosomal peptide synthases. Our ingestion feasibility gate also excluded or deferred over 50 candidate case studies, including whole-cell growth screens, coarse ordinal family screens, and libraries over 20,000 rows (such as the 160,000-variant four-site TrpB library), while staging recent imine reductase and carbene-transfer screens for future releases.

### Splits and noise estimation

The 30% similarity split removes close homologs but leaves distant ones (7.9 to 54.6% of test proteins have a training hit at 30% identity or more), and the 14 multi-substrate or multi-condition case studies are split by row, rewarding enzyme recognition over generalization (Section 3.5). Finally, the noise thresholds rest on two extra seeds (and four PLMs for attention pooling, EC number, and residue tasks) under a normal-noise assumption that slightly underestimates large seed gaps (Appendix E), and the residue tasks select epochs on validation loss rather than AUPRC due to memory cost.

## K Implementation and Community Leaderboard

Enzymado is implemented in Python with PyTorch and XGBoost. The PLMs run only in inference mode, and the heads are trained on precomputed embeddings stored as bfloat16 safetensors shards keyed by sequence, totaling 4.6 TB for the 14 PLMs. Special tokens are stripped so that position *i* of an embedding corresponds to residue *i*. Because models differ in whether they add start or end tokens, and a one-position shift would be invisible in a pooled task but would corrupt every residue label, we check the alignment for each model. Every run writes a manifest alongside its predictions that records a content hash of the dataset file, an inventory hash of the embedding store, hashes of the source tree and lockfile, the resolved package versions, the random seed, and whether deterministic kernels were enabled, so two runs that claim to be the same experiment can be checked against each other. The manifest also prevents a checkpoint from resuming in a changed environment, where the result would match neither the old experiment nor the new one.

Enzymado provides the datasets, cleaning protocols, split definitions, evaluation scripts, and model comparison outputs, along with a visual leaderboard for a single property, a task category, a split protocol, or the global rating. Adding a representation does not require rerunning the benchmark: because the heads, splits, and evaluation code are fixed, a new entry only needs embeddings for the shared sequence index and the five baselines trained on them before it joins the paired comparisons. We publish per-sample predictions along with the summary scores, so readers who disagree with our metrics, weights, or intervals can recompute them without the models or compute that produced them.

## L Hardware and Computational Requirements

Each job ran on one NVIDIA L40S GPU on nodes with Intel Xeon Gold 6338 CPUs and 4 to 32 cores per job, except for 141 runs (1.8%) on Intel Xeon Platinum 8570 nodes in a second cluster. All heads train on the GPU except the residue XGBoost heads, which train on the CPU with 4 threads and an on-disk cache of their binned features. The software stack consisted of Python 3.13, PyTorch 2.7.0 with CUDA 12.6, and XGBoost 3.2.0. Embeddings are extracted once and stored in bfloat16, requiring 4.6 TB for per-residue shards and 11 GB for mean-pooled vectors across the 14 PLMs. Mean-pooled heads cast them to 32-bit floats, whereas attention pooling runs its forward pass in bfloat16 and computes the loss in 32-bit floats. Table 14 reports the time and memory for each kind of run. The 4,200 benchmark runs took about 1,800 GPU-hours and the 3,844 extra seed runs about 1,050, nearly all of it spent on EC number with attention and on the residue tasks, while the scalar tasks with mean pooling train in seconds. These times cover training and prediction for the final runs, but not embedding extraction, the bootstrap, or failed attempts. Memory is the amount requested per job; resident memory reaches this amount because the embedding stores are memory-mapped.

**Table 14:** Compute per kind of run: median wall time per run in minutes for each head, total hours over the 8,044 runs (benchmark and extra seeds), and memory requested per job.

| **Tasks** | **Pooling** | **Linear** | **MLP** | **XGBoost** | **Total h** | **Memory (GB)** |
|---|---|---|---|---|---|---|
| scalar property | mean | 0.1 | 0.1 | <0.1 | 2 | 48 |
|  | attention | 8.1 | 8.4 | – | 166 | 124 |
| EC number | mean | 12.0 | 10.2 | 23.6 | 36 | 256 |
|  | attention | 294.4 | 466.5 | – | 842 | 124 |
| residue | mean | 161.2 | 236.8 | 128.6<sup>*</sup> | 886 | 160–256 |
|  | attention | 199.5 | 203.9 | – | 743 | 124 |
| case studies | mean | <0.1 | 0.1 | <0.1 | 5 | 48 |
|  | attention | 4.7 | 4.1 | – | 163 | 48 |

<sup>\*</sup>On CPU.

## M Cards and Dataset Ingestion

### M.1 Benchmark, Dataset, and Model Cards

#### Benchmark card

benchmark_card.json, at the repository root, has four blocks:

- the task registry, with each task’s type, scope, columns, deduplication keys, and split files;
- shared definitions referenced by every dataset card, covering cleaning, split protocols, standardization on the training split, and the two MMseqs2 settings (30% identity and 80% coverage for similarity splits, 50% identity and 80% coverage for the redundancy statistic);
- global statistics recomputed from disk;
- the rating configuration of Section 2.5.

A repository check, `scripts/ci/check_registry_card_parity.py`, fails if the task registry in the code and the benchmark card disagree.

#### Dataset cards

The computed card is written by a script from the processed and split files, and generation stops if a required tool or file is missing. It reports:

- label summaries by label kind: moments for regression, prevalence for binary tasks, annotated-residue density for residue tasks, and class counts per EC level;
- sequence lengths per row, per distinct protein, and over the sequences every PLM embedded;
- the ratio of MMseqs2 clusters to sequences at 50% identity, which ranges from 0.0002–0.03 in single-parent libraries through 0.15 in EC number to 0.66 in *k*_cat_.

The enriched card refers to these statistics rather than copying them. It is organized in six sections: how to think about the dataset (the observable, the effective sample size, and the claims it supports); label noise and the ceiling it implies; specificity and breadth; the experimental measurement, following STRENDA [68]; how the dataset was built, with a reconciliation from source to processed rows; and what each split tests.

#### Model cards

Model cards follow [54] and dataset cards follow [27]. Each model card describes the PLM along four axes:

- pretraining objective;
- pretraining corpus and its curation;
- context length and positional encoding;
- embedding width, regularization, and numerical precision.

It also documents our extraction code: which special tokens are removed, that ProstT5 receives its `<AA2FOLD>` prefix, that Ankh3 receives no `[NLU]` or [S2S] prefix, that ESM3 receives sequence only with its structure and function tracks left empty, and that E1 runs without retrieved homologs. An automated scan confirms that no Enzymado rating, rank, or score appears in any model card.

#### Quote verification

Each publication-derived field stores a value, a status (reported, interpreted, not reported, estimated, code, or computed), a verbatim quote of at most 250 characters, the source (main text or supplement), and the 1-based page. The verifier extracts the text of the cited page with pdftotext, both with and without layout mode, normalizes ligatures, dashes, quotation marks, and whitespace, and requires an exact substring match. Any card with a missing quote, a missing schema key, or an invalid reference to its computed statistics fails the check. All 1,895 quotes pass: 939 in the 14 model cards and 956 in the 46 dataset cards. The check establishes that the quoted text exists on the cited page; whether the quote supports the stated claim is not checked automatically.

### M.2 Task Layouts

Every dataset is converted to one of four layouts (Table 15), which fix the columns the loader expects and how features are built (Section 2.3).

**Table 15:** Canonical task layouts.

| Layout | Required columns | Features | Examples |
|---|---|---|---|
| A, protein | sequence, target | pooled embedding (+ conditions) | pH<sub>opt</sub>, EC number, PETase |
| B, protein–ligand | sequence, SMILES, target | pooled embedding + 200 RDKit descriptors | *k*<sub>cat</sub>, *K*<sub>M</sub>, binding |
| C, mutation | sequence, wild type, target | mutant minus wild-type embedding | Δ*T*<sub>m</sub>, ΔΔ*G* |
| D, residue | sequence, per-residue labels | *L* × *D* embedding | active sites, binding sites |

### M.3 Ingestion Workflow

The workflow is specified in `docs/AGENTIC_INGESTION_WORKFLOW.md` and `add_data.md`, and can be followed by a person or by a coding agent.

1. **Parsing.** Extract wild-type sequences, apply mutation codes, canonicalize substrate SMILES with RDKit, and record assay conditions.
2. **Feasibility gate.** Require a readout attributable to the enzyme under study, 300 to 20,000 rows (the 106-row *β*-glucosidase melting set of the Protein Engineering Tournament predates this check), at least 10 distinct sequences, and no overlap with the property datasets. Direct catalytic and biophysical assays qualify, and so do selections in which the enzyme’s own reaction is the selected step, such as the AmiE amidase and TrpB growth scans; whole-cell screens whose readout depends on unrelated cellular processes do not.
3. **Conversion.** A deterministic script per study maps the data to a layout and merges replicates by sequence, substrate, and conditions. Targets spanning two or more orders of magnitude are converted to log_10_ with the offsets of Appendix H.
4. **Sequence index.** New sequences are normalized and deduplicated against the shared pool, then appended with task-prefixed identifiers. Existing identifiers are never renumbered, so stored embeddings remain valid and only new sequences are embedded.
5. **Splitting.** Case studies use the random split by row, stratified on the target (Section 2.2). Regression targets are standardized on the training split.
6. **Registration.** The task is added to the code registry and to the benchmark card, the parity check is run, and both dataset cards are generated.
7. **Embedding.** The 14 PLMs embed the new sequences.
8. **Training and evaluation.** The five architectures are trained, and metrics and the 1,000 paired bootstrap resamples are computed.
9. **Rating.** The case-study leaderboard is refitted.

The workflow lists recoveries for the failures we met, such as dataset identifiers colliding with existing sequence identifiers, sequences missing from the index, and registry mismatches.

Because ratings are relative to the pool of datasets and competitors, adding a dataset changes the case-study ratings. Each leaderboard file therefore records the datasets it was fitted on.

### M.4 Curation Findings

Following the workflow, the agent surfaced problems in source data that would otherwise have entered training silently.

#### PET hydrolases

The source table [22] measures 19 enzymes at five temperatures and six buffer and pH conditions, for 570 rows. One row was labeled “30*^◦^*C, C6” a second time, directly before the 40 *^◦^*C block, which lacked its C6 row; its values lie between those at 30 and 50 *^◦^*C. The conversion script relabels it as 40 *^◦^*C, C6, and the dataset card records the correction.

#### Galactose oxidase variants

The source text [65] reports 1,325 variant–substrate measurements, whereas the deposited table holds 1,229 rows. Two pairs of enantiomers, (R)/(S)-1-phenylbutan-1-ol and (R)/(S)-2-chloro-1-phenylethan-1-ol, share one chiral SMILES string each in the deposited substrate file. Merging on sequence and SMILES therefore leaves 1,183 rows and 54 substrates. The 2D RDKit descriptors used for substrates do not distinguish enantiomers in any case, so this task cannot reward stereoselectivity.

#### UniProt fragments

Database exports include sequences that UniProt flags as fragments. Removing them (Section 2.2) drops 5,443 rows under the random split, 3,357 of them from EC number, and each dataset card reconciles the removed rows.

### M.5 Card-Generated Hypotheses and Empirical Tests

Because the model cards contain no Enzymado results, pairing a model card field with a dataset card field defines a testable hypothesis before looking at the leaderboard. Below, we describe the card fields, rationale, and results across the 56 task-splits (10 property tasks across 2 splits plus 36 case studies) for the five hypotheses in Section 3.6.

#### *H*_1_: Intra-family scaling and substrate feature dilution

We pair the model card fields `family, size_variant`, and `d_model` with the dataset card fields layout and `has_ligand_smiles`. Five PLM families have a smaller and a larger checkpoint trained with the same objective and corpus: ESM2 (650M at *D* = 1,280 to 3B at *D* = 2,560), Ankh (base at *D* = 768 to large at *D* = 1,536), Ankh3 (large at *D* = 1,536 to XL at *D* = 2,560), ESM-C (300M at *D* = 960 to 600M at *D* = 1,152), and Profluent E1 (300M at *D* = 1,024 to 600M at *D* = 1,280). Across the 56 task-splits, 37 take only the protein sequence (`has_ligand_smiles` = false, Layouts A, C, and D), whereas 19 append the 200 RDKit substrate descriptors to the *D*-dimensional protein embedding (`has_ligand_smiles` = true, Layout B: *k*_cat_, *K*_M_, binding, and 13 case studies). For linear and MLP heads, doubling the protein embedding width while keeping the 200 substrate descriptors fixed shrinks the substrate’s share of the input features from 20.7% (200/968) to 7.2% (200/2,760). This feature dilution should weaken or reverse the benefit of scaling within a family for linear and MLP heads on Layout B, while leaving decision trees (XGBoost) unaffected. With the mean-pooled MLP head, the larger model beats the smaller one on 30 of the 37 protein-only splits (81.1%, median gain +0.0091), but on only 3 of the 19 Layout B splits (15.8%, median change 0.0102). This reversal holds within each family on protein-only versus Layout B splits: Ankh3 wins 28/37 versus 3/19, ESM2 wins 24/37 versus 5/19, E1 wins 24/37 versus 3/19, and Ankh wins 25/37 versus 6/19. The mean-pooled linear head shows the same pattern, with the larger model winning 30/37 protein-only splits (81.1%, median +0.0323) versus 4/19 Layout B splits (21.1%, median *−*0.0038). In contrast, XGBoost gains from scaling on protein-only splits (31/37 wins, 83.8%, median +0.0096) and is unaffected on Layout B (9/19 wins, 47.4%, median *−*0.0008).

#### *H*_2_: Embedding width and attention-pooling overfitting

We pair the model card embedding width `d_model` (*D* [768, 2,560]) with the attention-pooling heads across both leaderboards. Mean pooling adds no parameters, whereas attention pooling learns query, key, value, and output projections on the *D*-dimensional residue vectors. Because its parameter count grows as (*D*^2^), an attention block has 11.1 times more parameters per residue at *D* = 2,560 (ESM2 3B and Ankh3-XL) than at *D* = 768 (Ankh-Base), which should cause wider embeddings to overfit. Across all 14 PLMs, `d_model` correlates negatively with the rating change from adding attention (rating_attn_ rating_mean_). On the property leaderboard, Spearman *ρ* with `d_model` is 0.84 for the MLP head, 0.81 for the linear head, and 0.91 for their average; on the case-study leaderboard, the correlations are 0.85, 0.79, and 0.86. At *D* = 768 (Ankh-Base), adding attention raises the linear head rating by +69.0 points (1469.4 to 1538.4) and the MLP rating by +3.7 points (1645.8 to 1649.5, second overall on the property leaderboard); at *D* = 960 (ESM-C 300M), it raises the MLP rating by +12.0 points on properties and +21.6 points on case studies. At *D* = 2,560 (ESM2 3B and Ankh3-XL), adding attention to the MLP drops property ratings by 189.6 and 198.6 points and case-study ratings by 484.1 and 428.9 points, placing the *D* = 2,560 attention models in the last four PLM spots on both leaderboards.

#### *H*_3_: Structural pretraining and expression decoupling

We pair the model card fields `base_model` and `pretraining_objectives` for ProstT5 and ProtT5-XL with the dataset card `field is_expression_decoupled`. ProstT5 starts from ProtT5-XL (with the same 24-layer, *D* = 1,024 architecture) and is trained further to translate between amino acid sequences and Foldseek 3Di structural tokens. In the dataset cards, 28 datasets (37 task-splits) measure purified or concentration-normalized properties (`is_expression_decoupled = true`: *k*_cat_, *K*_M_, pH_opt_, Δ*T*_m_, ΔΔ*G*, active and binding sites, EC number, and purified enzyme panels), whereas 18 datasets (19 task-splits) measure readouts that depend on cellular abundance or heterologous expression (`is_expression_decoupled = false`: DeepSol solubility, split-GFP solubility, VAMP-seq abundance, unnormalized cell-lysate screens, and growth-coupled deep mutational scans). Training on 3Di tokens emphasizes 3D backbone geometry over the surface composition and translation signals that govern expression and aggregation, so ProstT5 should beat ProtT5-XL on expression-decoupled tasks and lose to it on expression-confounded tasks. With the mean-pooled MLP head, ProstT5’s largest gains over ProtT5-XL all occur on expression-decoupled tasks: EC number (+0.103 AUPRC on the similarity split, 0.826 versus 0.723), Δ*T*_m_ (+0.063 *τ* on similarity, 0.326 versus 0.263), pH_opt_ (+0.040 and +0.022 *τ* on random and similarity), active sites (+0.039 AUPRC on similarity, 0.946 versus 0.907), *K*_M_ (+0.031 *τ* on similarity), and the purified copper radical oxidase panel (+0.027 *τ*). Conversely, its largest losses to ProtT5-XL occur on expression-confounded tasks: DeepSol solubility (0.138 and 0.080 AUPRC on similarity and random splits, 0.661 versus 0.799 and 0.783 versus 0.864), amidase growth selections on propionamide (0.065 *τ*) and acetamide (0.054 *τ*), and split-GFP solubility (0.055 *τ*). Overall, ProstT5’s mean rank among the 14 PLMs improves from 7.74 on expression-confounded tasks to 5.64 on expression-decoupled tasks.

#### *H*_4_: Protein multiplicity and sequence memorization

From the computed dataset card statistics (`rows` and `distinct_proteins`) and split rules (`grouping_unit_and_what_it_tests`), we define protein multiplicity as *M* = rows/distinct_proteins. Across the 46 datasets, *M* = 1.0 when every row is a distinct sequence (the 22 single-enzyme variant libraries and 3 property datasets: active sites, binding sites, and solubility), and *M >* 1.0 whenever a sequence keeps several rows, either from repeated entries or measurements (1.0002 in EC number, 1.03 in pH_opt_, 1.44 in Δ*T*_m_, and 1.49 in ΔΔ*G*) or because the same enzyme is measured on several substrates or conditions (from 2.5 in binding and 3.3 in *K*_M_ up to 30.0 in PETase, 34.8 in galactose oxidase, and 96.0 in the esterase panel). As *M* grows, and especially under row-stratified case-study splits where the same sequence appears in training and test sets, the task shifts from generalizing to new sequences to recognizing a known enzyme and splitting on the substrate or condition features. Across all 56 task-splits, *M* correlates positively with the lead of XGBoost over the MLP (Spearman *ρ* = +0.73; within the 36 case studies, *ρ* = +0.74) and negatively with the lead of the best PLM over the best no-plm baseline (*ρ* = 0.73; within case studies, *ρ* = 0.79). Splitting the 36 case studies by *M* also explains where the two leaderboards disagree: on the 22 single-enzyme variant libraries (*M* = 1.0), where every test sequence is a new mutant, the T5 span-denoising models take the top four mean ranks across tasks (Ankh-Large at 4.61, Ankh-Base at 4.80, ProtT5-XL half at 4.95, and ProtT5-XL at 5.02, ahead of Ankh3-XL at 6.16, ProstT5 at 7.07, ESM2 3B at 8.07, and ESM3 at 8.73), whereas on the 14 multi-substrate or multi-condition panels (*M >* 1.0), ProstT5 (6.04), ESM2 3B (6.57), and ESM3 (6.71) rank highest and XGBoost beats the MLP on 13 of the 14 datasets.

#### *H*_5_: Assay noise and STRENDA completeness on kinetic tasks

We pair the dataset card fields `biochemical_observable, noise_regime`, and `strenda_conditions` (which records whether temperature, pH, buffer composition, ionic strength, cofactors, substrate concentration, and reaction time are controlled) across matched kinetic tasks. In the property suite, *k*_cat_ and *K*_M_ from CatPred-DB have noise_regime = `inter_lab_aggregated` and 0 of 7 STRENDA conditions controlled or given to the model: their labels combine BRENDA and SABIO-RK measurements from many studies under unrecorded conditions, with *k*_cat_ keeping the maximum reported value per enzyme–substrate pair (an estimated inter-lab MAE floor of 0.38 to 0.71 log_10_ units). In the case-study suite, the PafA alkaline phosphatase datasets (`pafa_htmek_kcat, pafa_htmek_km`, and `pafa_htmek_kcat_km`) measure the same Michaelis–Menten constants under `low_noise_multi_replicate_microfluidic` conditions with all 7 of 7 STRENDA fields fixed on one microfluidic platform (10 to 15 replicates per variant, 95% confidence interval 0.07 log units). If the small gap between PLMs and no-plm on database kinetics comes from unmodeled assay noise rather than a lack of kinetic information in the embeddings, then holding the assay conditions fixed should widen that gap. On database *k*_cat_ (random split), the best PLM scores *τ* = 0.388 against 0.356 for no-plm, a gain of only +0.031 (and on database *K*_M_, *τ* = 0.515 against 0.488, a gain of +0.028). On microfluidic PafA *k*_cat_, the best PLM scores *τ* = 0.479 against 0.324 for no-plm, a gain of +0.155 (5.0 times larger than on database *k*_cat_). On microfluidic *k*_cat_/*K*_M_, the PLM lead over no-plm grows 8.5-fold to +0.266 (*τ* = 0.423 against 0.157), and on microfluidic *K*_M_ it grows 2.8-fold to +0.077. Across all 46 regression task-splits, datasets marked as `inter_lab_aggregated` (*n* = 8) reach a median best-PLM *τ* of 0.471 (mean 0.441), compared with 0.543 (mean 0.550) for single-lab controlled assays (*n* = 38).

## N How to Use Enzymado

Enzymado is open-source on GitHub (Appendix Q) and is set up to be straightforward to reproduce and extend. Users can reproduce it at three levels: the property and case-study leaderboards (level one), a single task (level two), or the entire benchmark of 4,200 runs (level three). Detailed instructions and terminal commands are available in REPRODUCE.md.

At level one, users generate the leaderboards from the pipeline results stored in the results folder of the main branch. At level two, users compute PLM embeddings and train prediction heads on an already-split dataset from data/processed/, and then compare their results against the published intervals. At level three, users re-run the entire benchmark on a Slurm GPU cluster from the shared sequence pool and the processed datasets. Levels one and two take minutes to hours, whereas level three may take several days because it re-embeds all PLMs, retrains all heads, and refits the leaderboards.

Users can also add their own datasets and predictor heads and then re-run parts of the benchmark on them. New datasets are added as a CSV file in one of three layouts: protein only, protein with substrate, or mutant with wild type. Their sequences are split randomly or by sequence similarity, then embedded, trained, and scored with the same 95% bootstrap intervals. New predictor heads are added as a PyTorch nn.Module in models/nn_blocks.py. Detailed instructions for both are available in add_data.md. New heads receive their own metrics and intervals, but they do not change the published leaderboard.

## O Ethical and Data Licensing Statement

Each source dataset used in Enzymado has its own license terms. Users should consult the original providers (BioLiP, CatPred-DB, BRENDA, FireProtDB, UniProt/Swiss-Prot, Proteinea, and the authors of the case-study datasets) for the applicable licenses before commercial use or redistribution.

## P Generative AI Use Disclosure

This work implemented generative AI for code and website development, as well as for writing support.

## Q Resource Availability

- >The Enzymado leaderboard with LLM integration is publicly available on the Enzymado official website at https://huggingface.co/spaces/Enzymado/Enzymado.
- >Source code and analysis scripts are available on GitHub at github.com/chemcognition-lab/enzymado-benchmark.

